# Multimodal measures of spontaneous brain activity reveals both common and divergent patterns of cortical functional organization

**DOI:** 10.1101/2022.12.13.520297

**Authors:** Hadi Vafaii, Francesca Mandino, Gabriel Desrosiers-Grégoire, David O’Connor, Xilin Shen, Xinxin Ge, Peter Herman, Fahmeed Hyder, Xenophon Papademetris, Mallar Chakravarty, Michael C. Crair, R. Todd Constable, Evelyn MR. Lake, Luiz Pessoa

## Abstract

Work in humans and animals shows that the brain can be decomposed into large-scale functional networks. Whereas most studies, especially in humans, use the blood-oxygenation-level-dependent (BOLD) signal, the relationship between BOLD and neuronal activity is complex and incompletely understood. This limits our ability to interpret and apply measures derived from fMRI-BOLD. Here, we employ wide-field Ca^2+^ imaging simultaneously recorded with fMRI-BOLD in highly-sampled mice expressing GCaMP6f in excitatory neurons. These unique data enabled us to characterize the similarities and differences between networks discoverable by each modality. Importantly, we applied a network partitioning approach that uses a mixed-membership algorithm, which allows brain regions to participate in multiple networks with varying strengths. This contrasts with assuming regions belong to only one network. Our findings demonstrate that (1) most BOLD networks are detected via Ca^2+^ signals. (2) There is considerable overlapping—as opposed to disjoint—network organization that is evident from both modalities. (3) Large-scale networks determined by Ca^2+^ signals at low temporal frequencies (0.01 – 0.5 *Hz*)—as opposed to higher frequencies (0.5 – 5 *Hz*)—are more similar to those determined by BOLD. (4) Despite many similarities, differences emerge across modes including the spatial distribution of membership diversity (the extent to which regions affiliate with multiple networks). In sum, Ca^2+^ imaging of excitatory neurons confirms that the mouse cortex is functionally organized into overlapping large-scale networks in a manner that reflects many, but not all, properties observable with simultaneous fMRI-BOLD; affirming the neural origins of patterns of brain organization that are evident in a clinically accessible neuroimaging modality.

## Introduction

Understanding the organization of large-scale brain networks remains a central problem in neuroscience, both in humans [1–7] and rodents [8–26]. Work in both species shows that the brain can be decomposed into several large-scale systems, such as the so-called “default network” [27, 28], using a variety of approaches. Understanding how brain networks support behavior, and are affected by injury and disease, holds great promise for future applications of these measures as diagnostic and prognostic tools. Yet, progress has been relatively slow partly because the bulk of what we know, especially in humans, is based on the blood-oxygenation level-dependent (BOLD) contrast obtained with functional magnetic resonance imaging (fMRI). The relationship between BOLD and neuronal activity is complex and incompletely understood [29–36], which has historically posed several challenges to interpreting networks discovered by this technique [37–39] and using them as clinical tools.

To address this gap, multimodal imaging in mice is emerging as a promising approach because a wide array of neuroimaging techniques are available for animal models (but inaccessible to humans) [40–43]. Despite the challenges of simultaneous multimodal acquisition, obtaining a more specific measure of neural activity, such as signals from fluorescent calcium-indicators (Ca^2+^) simultaneously with fMRI-BOLD (or another hemodynamic signal) enables the quantification and/or validation of the neural contribution to BOLD [40, 44–48]. Among reported multimodal studies, a few have evaluated the concordance across signal modalities of functional connectivity, a measure of inter-regional synchrony that is widely employed to characterize large-scale brain networks [40, 44, 49].

In particular, Murphy and colleagues compared functional connectivity results obtained with Ca^2+^ to those obtained with hemodynamic imaging [46]. The study reported that functional connectivity (i.e., pairwise correlations) exhibited good agreement between modalities, but approximately half of the pairwise correlations differed significantly. Further, previous work by us using simultaneous wide-field Ca^2+^ imaging and fMRI-BOLD showed that functional connectivity obtained with the two signal types showed good agreement overall, but notable differences were also detected [40]. Wide-field Ca^2+^ imaging is an emerging optical approach that captures a large field of view encompassing most of the mouse cortical mantle [50, 51] with relatively high resolution (e.g., 25 × 25 micron pixels). However, these prior studies investigated a rather coarse parcellation of the cortex (7 brain regions [46], and 10 brain regions [40] per hemisphere) and assumed, as most human and rodent research does, a disjoint functional organization.

Many complex networks, including biological, technological, and social ones, are inherently *overlapping* (nodes participate in multiple communities) rather than *disjoint* (each node belongs to a single community) [52–54]. In the brain, uncovering overlap would reveal that some regions participate in multiple networks, consistent with the notion that functionally flexible regions contribute to multiple brain processes [55–58]. Although evidence for overlap in human brain networks has accrued based on multiple analysis techniques [7, 58–60], it is unclear whether the putative overlapping organization is driven, at least in part, by the nature of BOLD signals.

Here, we use a simultaneously recorded wide-field Ca^2+^ and fMRI-BOLD dataset ([40]; Figure 1A) to resolve whether functional networks discovered with BOLD are also detected with Ca^2+^, while also determining their potential overlapping organization. This dataset contains a highly-sampled group of mice (10 animals, 3 sessions each, 4 per session; Figure 1B) expressing GCaMP6f in excitatory neurons, and it is substantially larger and of higher quality than our original work, which allowed us to use a much finer parcellation (~ 270 regions per hemisphere; Figure 1D). Further, these data are unique and constitute the second dataset of its kind (the first being our description of the method [40]).

**Figure 1:**
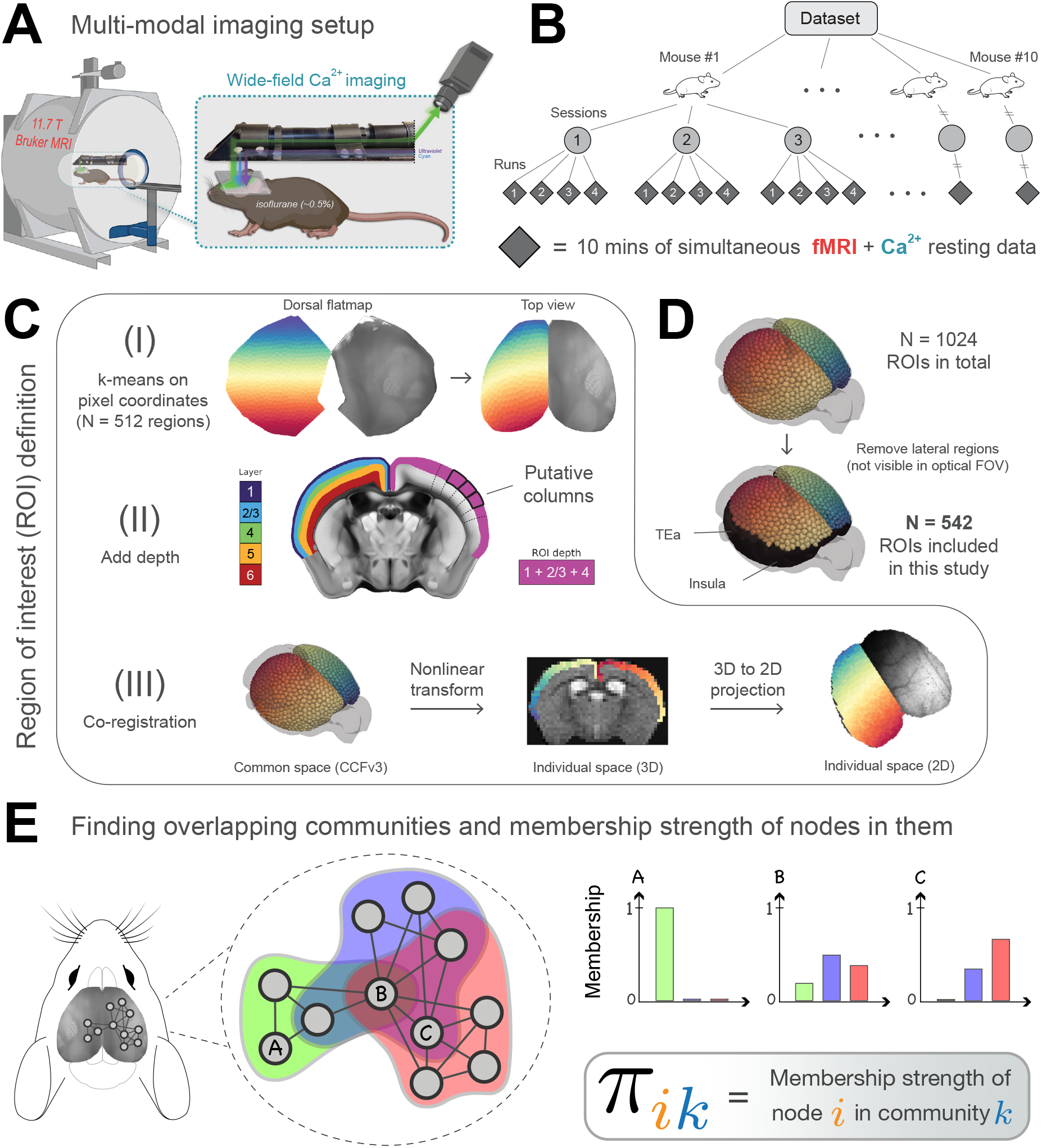
Experimental setup and overlapping community analysis. **(A)** Simultaneous fMRI-BOLD and wide-field Ca^2+^ imaging [40] on lightly anesthetized GCaMP6f mice. **(B)** Hierarchical structure of the data. *N* = 10 mice, scanned across 3 longitudinal sessions, with 4 runs per session each lasting 10 minutes. **(C)** Definition of regions of interest (ROIs) within the Allen Mouse Brain Common Coordinate Framework (CCFv3) space [70]. (I) Divide the mouse dorsal flatmap into a total of *N* = 1024 spatially homogeneous regions. (II) Add depth by following streamlines normal to the surface of the cortex. The resulting ROIs are “column-like”. (III) Transform ROIs from the common space into 3D and 2D individual spaces (Methods). Dorsal flatmap, layer masks, and columnar streamlines are available as part of CCFv3. **(D)** We restricted analysis to the subset of ROIs that appeared in the optical field of view (FOV) after co-registration (thus leaving out lateral areas such as the insula and temporal association areas). **(E)** We applied a mixed-membership stochastic blockmodel algorithm to estimate overlapping communities [61]. *Membership strength* (values between 0 and 1) quantifies the affiliation strength of a node in a network. Here, node *A* belongs only to the green community, node *B* belongs to all three communities with varying strengths, and node *C* belongs to the blue and red communities with varying strengths.

We investigated overlapping functional brain organization in multimodal mouse data by using a Bayesian generative algorithm that estimates the membership strength of a given brain region to all networks (Figure 1E) [58, 60, 61]. We report region-level properties which quantify various aspects of network organization such as degree [62, 63] and diversity [17, 64, 65], and contrast outcome measures obtained with fMRI-BOLD and Ca^2+^ signals. Further, given that the correlation of BOLD and Ca^2+^ signals is maximal when Ca^2+^ is temporally band-passed to match BOLD [40, 45], as well as the “lowpass” nature of the BOLD signal [66–68], we investigated whether measures were more closely matched within a slow (relative to fast) Ca^2+^ frequency range. More broadly, understanding whether and how network organization depends on the temporal frequency of the signals is important to help constrain the types of processes contributing to functional organization [69].

In sum, this work employed highly novel simultaneous wide-field Ca^2+^ and fMRI-BOLD to uncover the large-scale functional organization of the mouse cortex. Such advances are fundamental to advancing our understanding of brain organization and validating the neural origins of clinically accessible fMRI-BOLD measures.

## Results

Mice underwent multimodal imaging using an apparatus built for the simultaneous collection of wide-field Ca^2+^ and fMRI data, as described previously by us ([40]; see Methods). Head-fixed GCaMP6f mice were placed inside an MRI scanner into which we inserted optical components for Ca^2+^ signal acquisition (Figure 1A). Ten lightly anesthetized mice (0.5% isoflurane) were scanned across 3 longitudinal sessions; each session contained 4 runs, each lasting 10 minutes (Figure 1B).

We preprocessed BOLD data using the RABIES package [71], which has been applied to previous mouse [20] and rat [26] fMRI datasets. This included a simple high-pass filtering step [6] which resulted in a frequency range of 0.01 to 0.5 *Hz* for BOLD (acquisition rate was 1 *Hz*; see Methods). We considered Ca^2+^ data of two frequency bands: 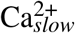 matching the BOLD frequency, and 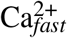 which was high-pass filtered between 0.5 and 5 *Hz* (effective acquisition rate was 10 *Hz*). Ca^2+^ data were processed as described by us previously [72].

To build functional networks, a common set of regions of interest (ROIs) was defined (Figure 1C; Methods). To relate 3D BOLD and 2D Ca^2+^ data, we adopted the CCFv3 space for the mouse brain provided by Allen Institute for Brain Sciences [70]. ROIs covered most of the cortex but left out lateral regions not well captured by wide-field Ca^2+^ imaging (Figure 1D). Correlation matrices were computed for each *run* by determining the pairwise Pearson correlation between ROI time series data and were binarized by keeping the top *d*% strongest edges. We present results mainly for 512 ROIs and *d* = 15% but explored other values to ascertain the robustness of the results across a range of parameter choices. We used a mixed-membership stochastic blockmodel [73] algorithm that generates overlapping networks [58, 60, 61]. The algorithm determines *membership* values for each brain region, with one value per network (Figure 1E). Membership can be thought of as a limited resource, such that they sum to 1, which allows membership values to be interpreted as probabilities. Overlapping networks were computed at the level of *individual runs*, such that membership values were determined for individual runs and averaged across sessions to determine an animal-level result. Random-effects group analysis was then evaluated based on animal-level estimates and variability.

### Determining overlapping network organization

We performed overlapping network detection as a function of the number of networks, across modalities. Recent work has shown decomposition of the mouse cortex into as few as 2-3 networks [17, 19, 74], but more typically has uncovered 7-10 networks [9, 14, 15, 20]. With three networks, our approach captured previously observed systems, namely, the visual (overlapping community 2, or OC-2) and somatomotor (OC-3) networks, as well as a large system (OC-1) that included cortical territories classified by some as the default network [13, 14, 18, 28] (Figure 2A) (we use the term *network* interchangeably with *overlapping community*). To facilitate comparison with standard algorithms, we also determined a disjoint version in which brain regions were assigned the network label with the largest membership value.

**Figure 2:**
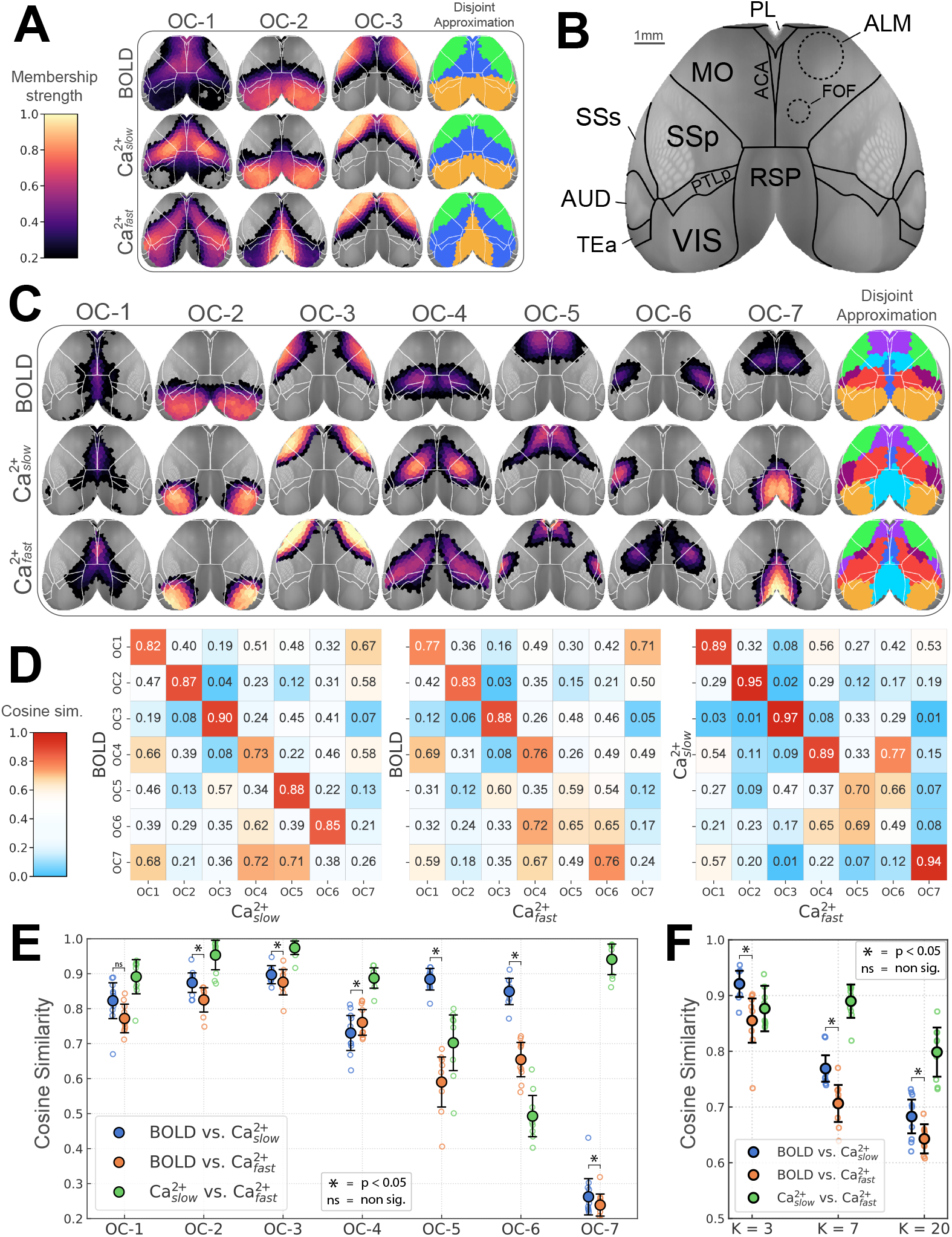
Overlapping functional networks of the mouse cortex. **(A)** Decomposition based on three networks. Color scale indicates membership strengths (see Figure 1E). The disjoint approximation is obtained by taking each node’s maximum membership value. OC, overlapping community. **(B)** Cortical areas (top view) with areas defined in Allen reference atlas [70]. In addition, dashed lines approximately correspond to functionally defined subregions in the secondary motor area [76, 80]. **(C)** Decomposition with seven networks. **(D)** Quantifying network similarity based on cosine similarity (1 = identical, 0.5 = “orthogonal”, 0 = perfectly dissimilar or “inversely correlated”). A custom color map was used to emphasize similarities and only strong dissimilarities. **(E)** Diagonal elements of matrices in D are shown. Empty circles correspond to individual animals; the large circle is the group average. **(F)** Overall similarity scores by combining all networks as a function of the number of networks (3, 7, and 20). **(E-F)** Comparison of BOLD and 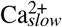 networks (*p* < 0.05, Holm-Bonferroni corrected). Error bars are 95% confidence intervals based on hierarchical bootstrap. Abbreviations: ACA, anterior cingulate area; ALM, anterior lateral motor cortex; FOF, frontal orienting field; MO, somatomotor areas; PL, prelimbic area; PTLp, posterior parietal association areas; RSP, retrosplenial area; SSp, primary somatosensory area; SSs, supplemental somatosensory area; VIS, visual areas. See also Figure S1, Figure S2, Figure S3, and Figure S4.

With seven networks, well-defined visual and somatomotor networks (OC-2 and OC-3, respectively) were again identified [15, 20], together with additional systems covering bilateral and well-defined cortical territories (Figure 2C). OC-1 encompassed medial areas including cingulate cortex but also extended more laterally. OC-4 spanned from medial to lateral areas, including somatosensory cortex; for both Ca^2+^ signals, it also included the frontal orienting field (FOF), a possible homolog of the frontal eye field in primates [75–79]. OC-5 largely overlapped with the anterior lateral motor area, a region involved in motor planning [80–83]; notably, for 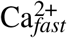 this network also included the supplemental somatosensory area. OC-6 overlapped with barrel field for BOLD and 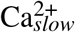, but captured upper limb somatosensory cortex for 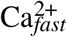. Finally, OC-7 was very different for BOLD and Ca^2+^ signals. For BOLD it was centered around FOF; for both Ca^2+^ signals, OC-7 was centered around the retrosplenial cortex. We also investigated a considerably larger number of networks (total of 20; Figure S1), revealing finer spatial networks that were mostly organized bilaterally. It is noteworthy that even with 20 networks, FOF did not appear as a separate network for either Ca^2+^ signals, in contrast to its appearance as such in the case of BOLD.

Next, we quantitatively compared the results of the different signals (BOLD, 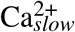, 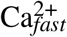) (Figure 2D). We focused on the subdivision of the cortex into seven networks, an intermediate number within the range frequently investigated in the literature [9, 14, 15, 20]. In particular, we evaluated the hypothesis that band-pass filtering Ca^2+^ signal to match that of BOLD signal leads to a Ca^2+^ network organization that more closely resembles that of BOLD. Networks were compared based on cosine similarity (a score of 1 indicates identical organization, 0.5 indicates “orthogonal/unrelated” organization, and 0 indicates perfectly “opposite” organization). The similarity between BOLD and 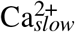 networks was relatively high (> 0.73), except for OC-7 (0.26), a network that was evident in both Ca^2+^ conditions but not captured by BOLD. In comparison, the similarity between BOLD and 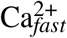 networks was generally lower but still relatively high for OC-1 to OC-4 (> 0.77), though modest for OC-5 and OC-6 (0.59 and 0.65, respectively). Combined, these results revealed that band-pass filtering Ca^2+^ signals had the strongest impact on the OC-5 and OC-6 networks (Figure 2E).

To further quantify the similarity between BOLD and Ca^2+^ network structure, we combined all networks to compute an overall index of similarity. In particular, we tested the prediction that the similarity between BOLD and 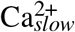 would be higher than that between BOLD and 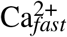 [40, 45, 49]. Indeed, this relationship was statistically significant when the cortex was partitioned into 3, 7, and 20 overlapping networks (*p* < 0.05, permutation test, Holm–Bonferroni corrected). Importantly, these results were stable across multiple processing parameters (Figure S3), including thresholding of the functional connectivity matrices, and the number of ROIs (Figure S2).

In summary, overlapping networks were reliably estimated from both BOLD and Ca^2+^ data. Although several properties of large-scale systems captured by BOLD were largely reproduced by Ca^2+^ data, notable differences were uncovered, too, and will be further explored below. Importantly, our analysis confirmed the hypothesis that BOLD networks are more similar to those obtained with Ca^2+^ data band-passed in the slow temporal frequency range of BOLD (but see Discussion). In what follows, we focused on the overlapping network organization with seven networks, which captures large-scale cortical systems of intermediate size, as most often investigated in the literature [9, 14, 15, 20]. Importantly, whereas the subsequent results are described at the group level, we confirmed that the basic organization of the seven networks was observed at the individual level (Figure S4). Thus, group-level properties, including network overlap, are not driven by the process of performing group analysis.

### Quantifying overlapping organization

To quantify overlapping organization, we examined the distribution of membership values across networks. Membership values range between 0 to 1, and sum to 1 across networks. Thus, if a specific region has high membership value for one of its networks, the total membership across the remaining networks will be low. In particular, a disjoint organization would be revealed if all regions have very high membership values for a single network (“right peaked” distribution; Figure 3A, left). In contrast, a roughly uniform distribution would correspond to a network whose nodes affiliate with multiple networks but with varying strengths (Figure 3A, middle); a more extreme form of overlap would be one in which nodes tend to not affiliate with any network very strongly (Figure 3A, right). To generate membership distributions, we considered the membership range (0.2,1.0]; values less than 0.2 were not considered so as to *conservatively* characterize network overlap. The last bin contained values larger than 0.8 based on our simulations with synthetic graphs with known ground truth [84]. Specifically, disjoint synthetic networks had all their membership values concentrated in the last bin (0.8,1.0] (see Figure S5).

**Figure 3:**
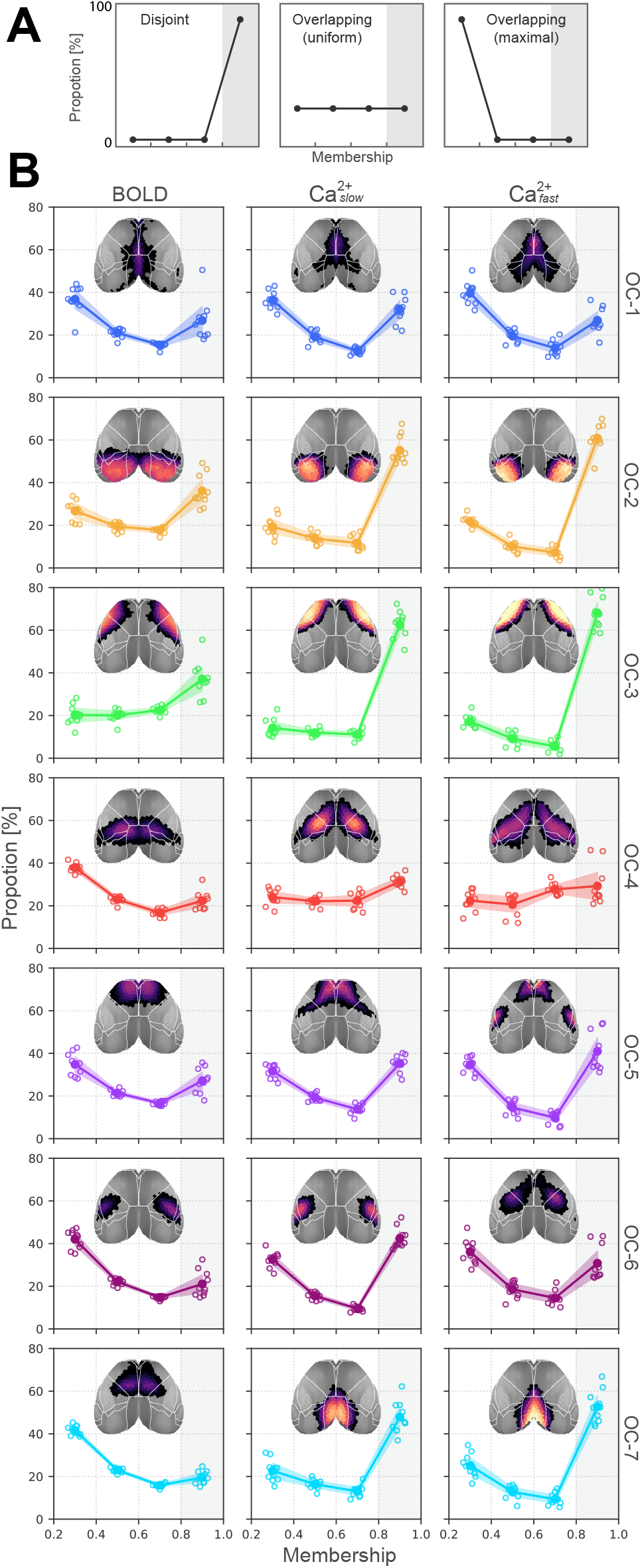
Distribution of membership values. **(A)** Three illustrative distributions. Left, disjoint organization with no small or mid-range memberships; Middle, overlapping with uniform weights in all membership values; Right, completely overlapping with no mid-range or stronger memberships. **(B)** Membership distributions computed from the real data hint at a substantial overlap. Note that here the y-axis is capped at 80%, indicating that none of the networks are truly disjoint. Empty circles correspond to individual animals; the large circle is the group average. OC, overlapping community. Compare with Figure S5 for membership distributions obtained for graphs with known ground truth overlap sizes.

Across conditions, most networks contained no more than 60% of brain regions with the highest level of membership strength (> 0.8); thus approximately 40% of the regions belonged to at least 2 networks, demonstrating substantial network overlap (Figure 3B). The least overlapping networks were the visual and somatomotor ones (OC-2 and OC-3) for BOLD, 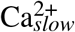, and 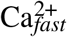, and the retrosplenial network (OC-7) for the two Ca^2+^ conditions. Among the most overlapping networks, across all conditions, OC-4 contained ~ 60% of the regions belonging to more than one network (see also OC-1). Taken together, our results demonstrate that the mouse cortex contains a substantially overlapping organization, which was present for BOLD and both Ca^2+^ frequency bands.

### Nested spatial organization of membership strengths

To further quantify network overlap, we considered four different threshold values and evaluated if a brain region’s membership strength statistically exceeded the values in question (FDR corrected; see Methods) (Figure 4A). Because there were seven networks, we considered thresholds that were multiples of 1/7. For example, regions in blue participated mostly in a single network (membership values > 3.5 × 1/7 = 0.5). In contrast, regions in yellow participated in more networks. Thus, regions in blue revealed territories more on the disjoint end of the spectrum, whereas those in yellow revealed territories with more overlap. Overall, overlapping network organization was arranged in a spatially coherent fashion that revealed spatial patterns of nested membership tiers.

**Figure 4:**
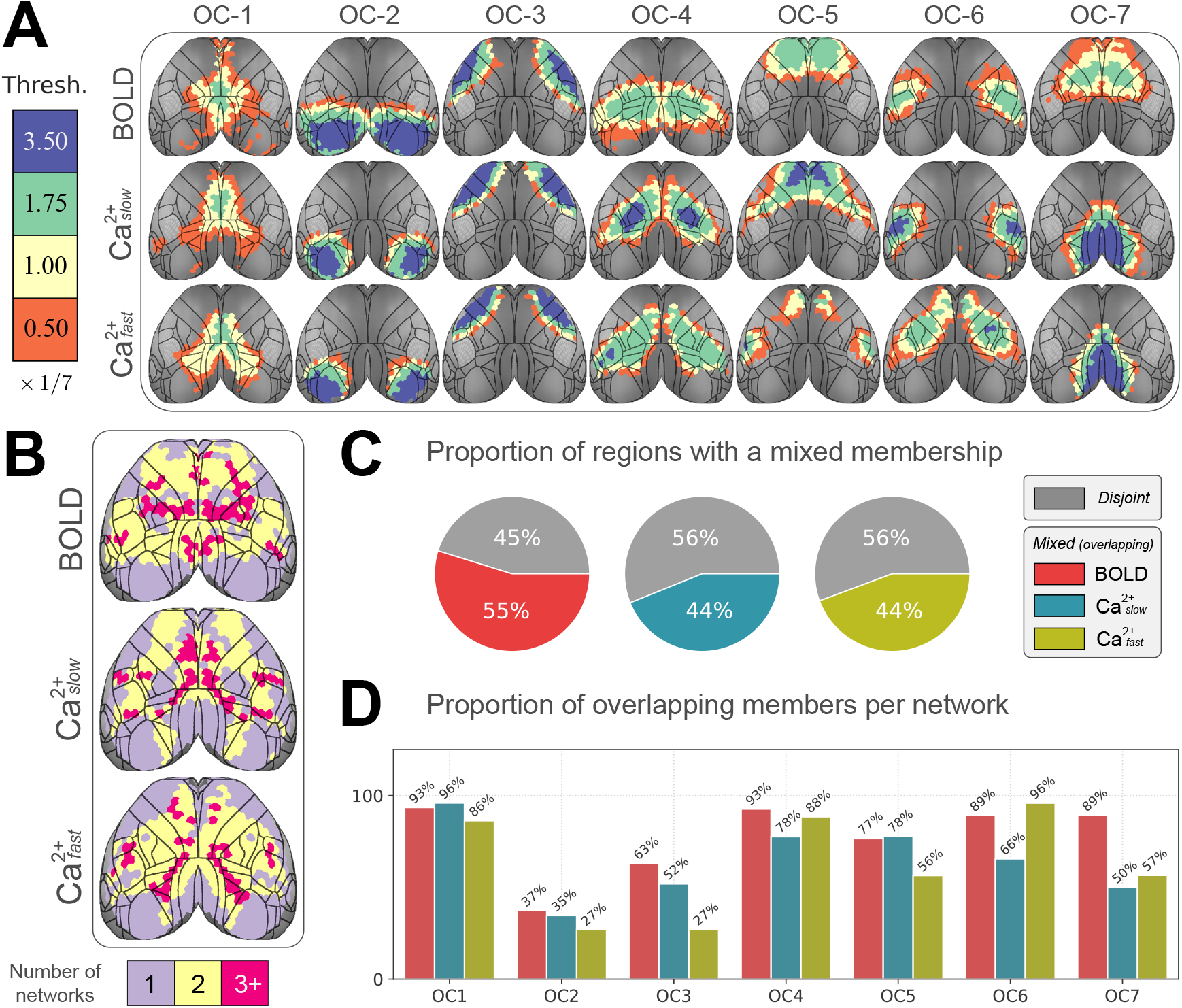
Quantifying overlap extent. **(A)** We used a statistical thresholding method to decide if a node *belongs* to a network. Members of networks are colored as a function of four thresholds. Blue (strong threshold, membership > 3.5 × 1/7 = 0.5) reveals faithful members, whereas orange (> 0.05 × 1/7) reveals the spatial extent of the network. Contour lines correspond to the fine division of the Allen reference atlas (see Figure S1B). OC, overlapping community. **(B)** Using a threshold of 1/7, we counted the number of networks that each brain region belongs to. Most regions belong to more than one network. **(C)** We defined a global *overlap score* for the entire graph as the ratio of overlapping regions divided by the total number of regions. **(D)** Overlap score at the network level is defined similarly. All networks are highly overlapping, except visual (OC-2) and somatomotor (OC-3) networks. A somewhat surprising finding is that for Ca^2+^, the retrosplenial network (OC-7) is relatively less overlapping than other networks.

### Majority of brain regions belonged to multiple networks

Here, we used the threshold approach of the previous section to quantify the number of networks an individual region “belongs to”. Specifically, if a region exhibited a membership value statistically greater than 1/7, the region was deemed to belong to the associated network (Figure 4B). By this definition, for both BOLD and Ca^2+^ signals, around half of the brain regions belonged to more than one network (Figure 4C).

To quantify network-level overlap, we determined the proportion of regions belonging to at least two networks. Network overlap was very high in multiple cases (Figure 4D). For BOLD, overlap was > 60% in all but one network; for 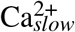, overlap was considerable and > 50% for all but one network; for 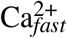 overlap was decreased but still > 40% in all but two networks (but note that OC-1, OC-4, and OC-6 exhibited scores > 85%).

In summary, approximately half of the brain areas of the mouse cortex belonged to at least two networks, a property observed with both BOLD and Ca^2+^ data. Simultaneously, particular large-scale cortical networks were highly overlapping, and even the least overlapping ones included more than a quarter of the regions that affiliated with more than one network.

### Membership diversity is spatially organized

The preceding analysis allowed us to estimate the extent of network overlap by utilizing a binary criterion for membership. However, such discretization discards continuous information that can be used to characterize a brain region’s *membership diversity* and its spatial arrangement across the cortex. To quantify such information here we computed the (normalized) Shannon entropy of the membership values of a region (Methods). A region that belongs to all communities with equal membership strengths will have maximum diversity; a region that belongs to a single network will have minimal diversity. In brief, membership diversity is potentially indicative of a region’s multi-functionality and/or involvement in multiple processes.

The distribution of membership diversity scores is shown in Figure 5A. A substantial number of regions exhibited diversity values very close to zero (for example, approximately 30% for 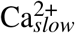), indicating that their membership strength was preponderantly associated with one network. Excluding this peak at zero diversity, the remaining nodes displayed values more or less along a continuum. However, of the regions associating with multiple networks, a considerable number of them affiliated with two communities (see the dashed line; e.g., a region with a membership value of 0.5 with one network, 0.5 with a second network, and 0 with the remaining networks).

**Figure 5:**
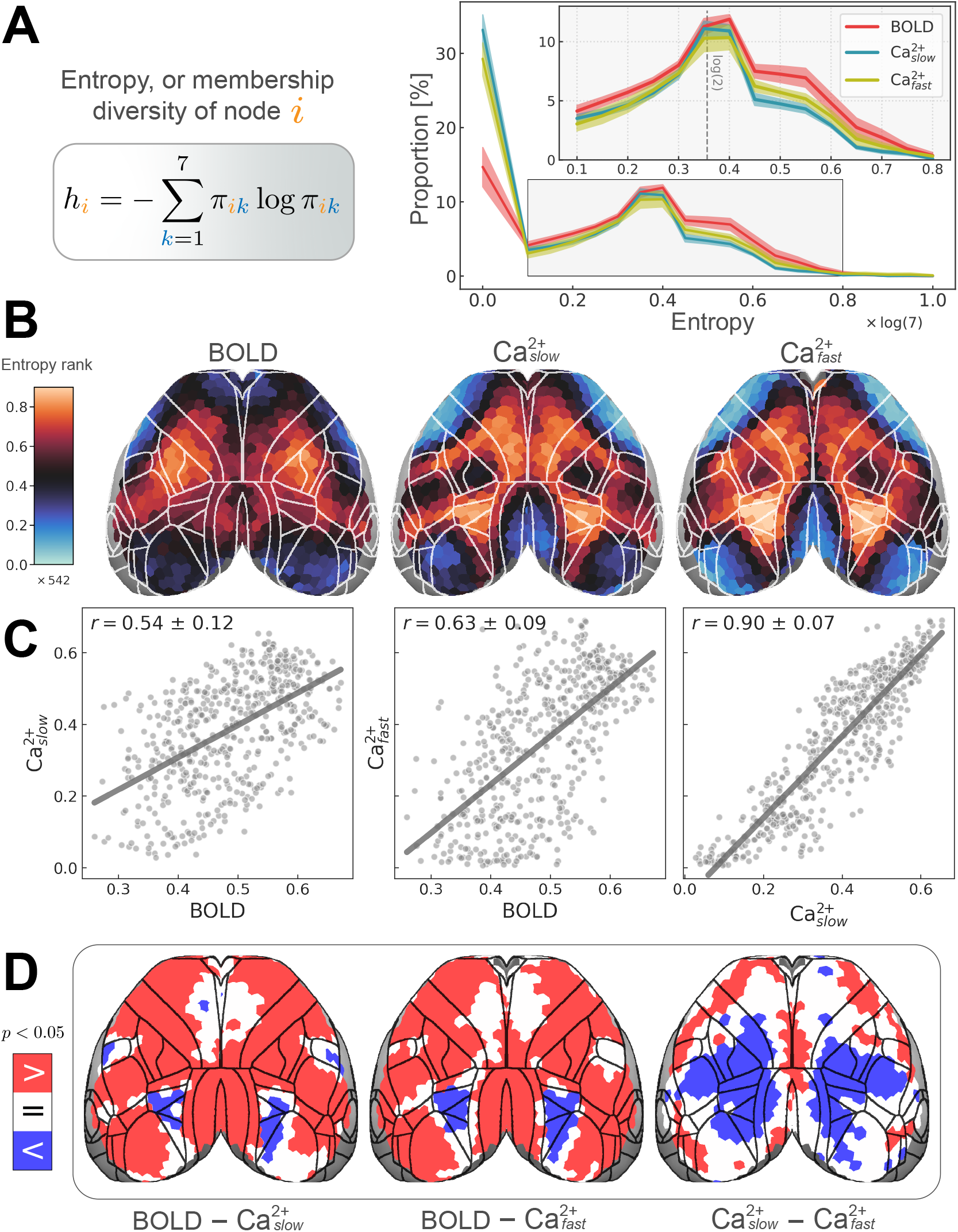
Introducing node entropy centrality. **(A)** Shannon entropy is applied to node membership vectors to quantify membership diversity. Values are normalized to the range [0,1]. *h_i_* = 0 if node *i* belongs to a single network (i.e., disjoint); *h_i_* = 1 if it belongs to all networks with equal strength (i.e., maximally overlapping). Right: distribution of node entropies. The largest peak occurs at *h* = 0, corresponding to disjoint nodes. The second largest peak occurs at *h* ≈ log(2) which corresponds to a situation where a node has a membership of 0.5 in two networks and 0 elsewhere. **(B)** Spatial distribution of region entropy. Rank-ordered to facilitate comparing across conditions (total of 542 ROIs); non-rank-ordered version in Figure S6A. Unimodal areas such as visual and somatomotor areas are low in entropy, whereas transmodal regions have higher entropy. See Figure S6 for a comparison with participation coefficient, a graph measure commonly used to quantify link diversity [17, 64, 65, 85, 86]. **(C)** Entropy values are positively correlated across modalities. **(D)** Permutation test revealed BOLD > Ca^2+^ in most regions, except for some frontal areas where BOLD = Ca^2+^, and higher visual areas where BOLD < Ca^2+^.

Next, we investigated the spatial distribution of membership diversity values. For visualization purposes, we rank-ordered values to inspect the overall pattern across conditions irrespective of absolute values; Figure 5B. For the non-rank-ordered version, see Figure S6A. The resulting patterns revealed partial agreement between BOLD and Ca^2+^ conditions, and especially strong agreement between the two Ca^2+^ frequency bands. To quantify the agreement between conditions, we (Pearson) correlated membership diversity values; between BOLD and 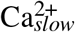: *r* = 0.54 ± 0.11; between BOLD and 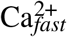: *r* = 0.63 ± 0.09; and between 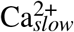 and 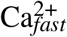: *r* = 0.90 ± 0.07 (variability obtained based on hierarchical bootstrapping; see Methods). Next, we explicitly compared membership diversity maps to determine ROIs that showed differences between conditions Figure 5B. We found that membership diversity was consistently larger for BOLD compared to both Ca^2+^ conditions, except for delimited sectors of the cortex. Intriguingly, 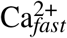 exhibited a large territory of ROIs with larger membership diversity compared to 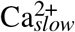.

In summary, overlapping networks can be described in terms of the diversity of their region’s affiliations. We found an intermediate amount of agreement in the spatial distribution of membership diversity across all conditions. We predicted that BOLD and 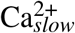 networks would be more similar compared to BOLD and 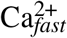. However, the effect was rather modest, except for two networks (OC-5 and OC-6).

### High degree nodes are substantially different across data modalities

In the preceding section, we compared network organizations in terms of membership diversity. Next, to further compare the results across the three conditions (BOLD, 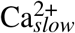, 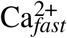), we investigated graph *degree*, another measure of node centrality that quantifies the number of functional connections of brain regions [62, 63]. The distribution of degree values for BOLD was shifted toward lower values compared to Ca^2+^, whereas the distributions for the latter were more similar to each other (Figure 6A). More importantly, the spatial distribution of degree differed considerably between BOLD and Ca^2+^ conditions (Figure 6B), which was reflected in the correlations across conditions (Figure 6C). Notably, degree values were highly correlated between the two Ca^2+^ conditions. Finally, to eliminate differences based solely on the magnitude/variability of the degree values, we obtained percentile maps by calculating t-statistics followed by rank-ordering [17] and found the same pattern of results (not shown).

**Figure 6:**
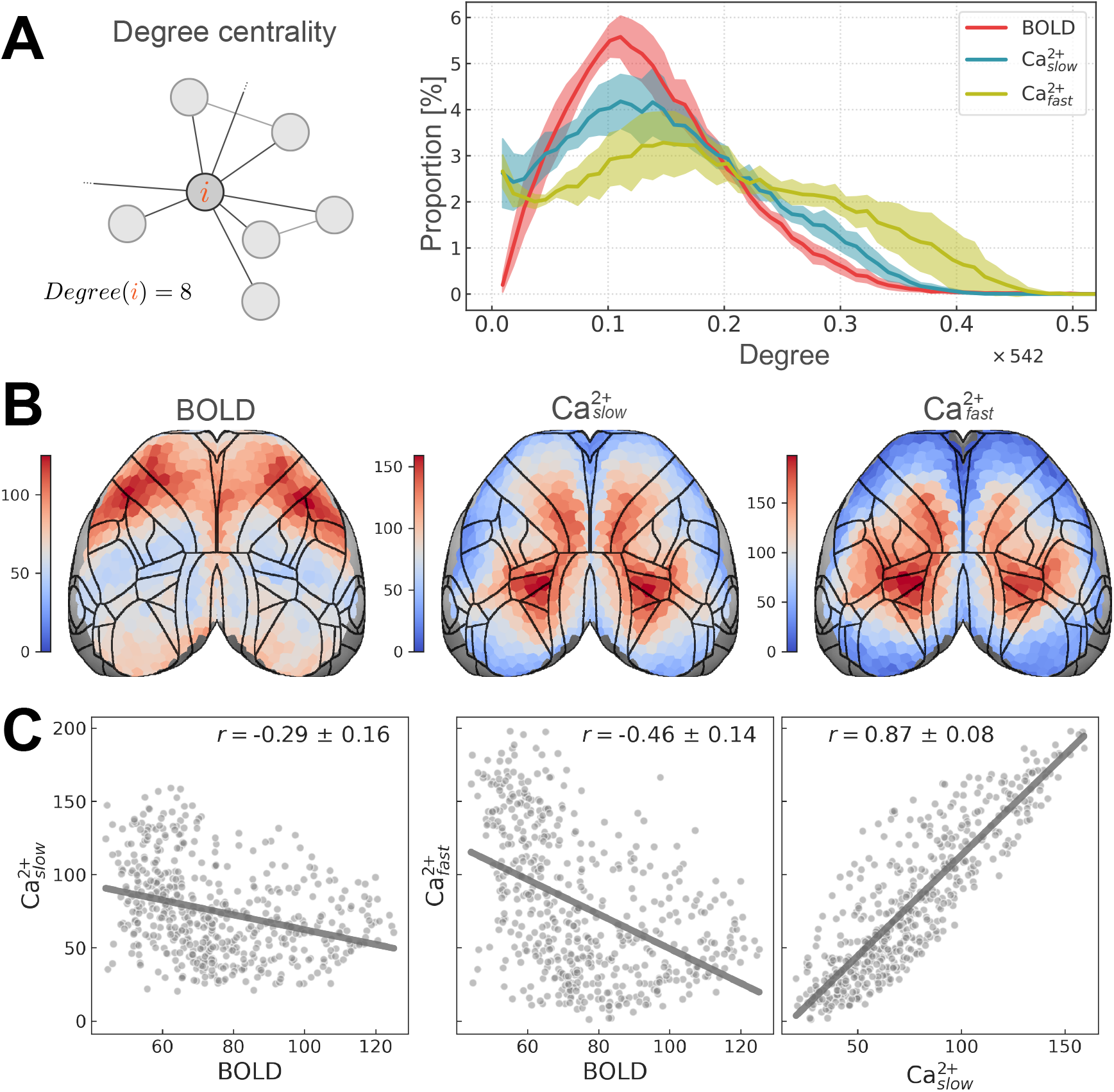
High degree nodes are substantially different across data modalities. **(A)** Node degree count is one of the most basic and widely used measures of network topology. Right: degree distributions are different across modalities, with BOLD being less dissimilar to 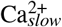 than 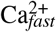. **(B)** Average node degrees visualized on the cortex reveals the discrepancy across modalities. **(C)** The dissimilarity is quantified as a negative (Pearson) correlation between BOLD and Ca^2+^. The discrepancy across modalities persisted for other choices of edge-filtering thresholds or preprocessing steps (see Figure S7).

### Network overlap also differs across modalities

Thus far, our findings support the existence of overlapping cortical networks, but we have not characterized the nature of this overlap. Here, we consider node degree and membership diversity (entropy) together, because their combination can further inform functional organization [85]. For example, network hubs can be classified as provincial or connector hubs based on their intra- and inter-community connectivity patterns [17, 86]. A possible scenario is one in which high-diversity regions have relatively low degree, a scenario we call *sparse* overlap (Figure 7A, left). Alternatively, high-diversity regions also have high degree, called *dense* overlap (Figure 7A, right). We visualized the pairwise entropy-degree relationship at the level of brain regions (Figure 7B). The two measures were inversely correlated for BOLD (r = −0.44 ± 0.16), more consistent with sparse overlap (high diversity nodes did not tend to have high degree). In contrast, entropy and degree were positively correlated for Ca^2+^ (*r* = 0.44 ± 0.09, 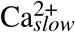; *r* = 0.69 ± 0.07, 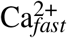), more consistent with dense overlap (high diversity nodes tended to have high degree).

**Figure 7:**
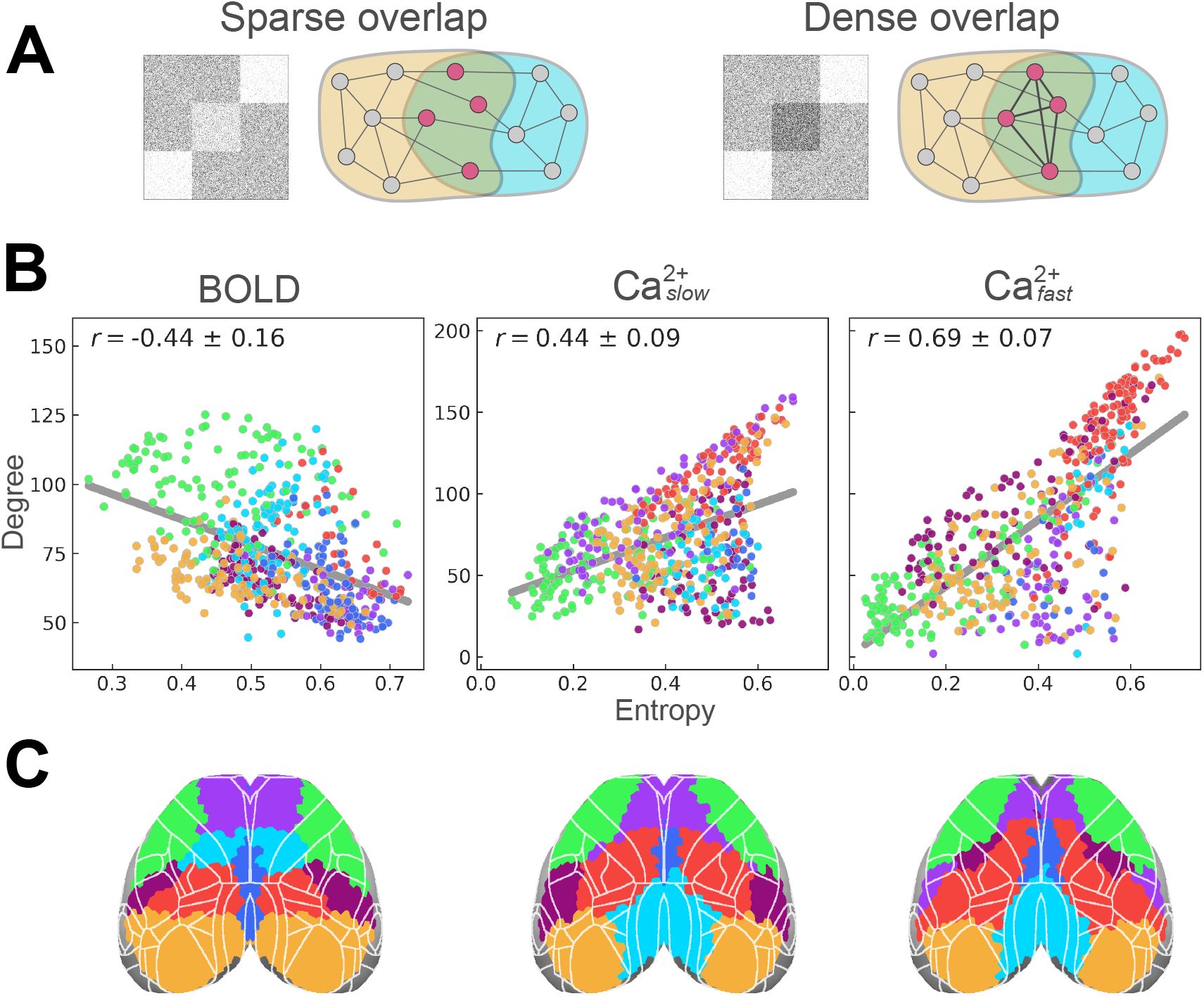
Network overlap also differs across modalities. **(A)** Possible types of overlap are illustrated. **(B)** Entropy (a measure of connection diversity) and degree (a measure of connection strength) are plotted. The negative correlation for BOLD indicates the presence of sparse overlap, reminiscent of previous reports in human fMRI [64, 65]. In contrast, the relationship is positive for both 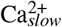 and 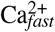, with a larger correlation for 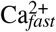. Thus, Ca^2+^ data is more consistent with dense overlap. **(C)** Nodes are colored according to their disjoint network assignment (see last column in Figure 2C). See also Figure S6.

## Discussion

We used highly novel simultaneous wide-field Ca^2+^ and fMRI-BOLD acquisition to characterize the functional architecture of the mouse cortex. The spatial organization of large-scale networks discovered by both modalities showed many similarities, with some temporal frequency dependence (BOLD networks were generally more similar to 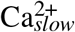 than 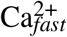). Functional connectivity interrogated using a mixed-membership algorithm, instead of traditional disjoint approaches, confirmed the hypothesis that mouse cortical networks exhibit substantial overlap when either BOLD or Ca^2+^ signals were considered. Further, despite the considerable agreement, we also uncovered important differences in organizational properties across signal modalities.

### Fine-grained functional networks of the mouse cortex revealed both inter-modal similarities and differences, with dependence on Ca^2+^ temporal frequency

Previous multimodal studies comparing cortical functional organization via concurrent GCaMP6 Ca^2+^ and hemoglobin-sensitive imaging have predominantly employed seed-based analyses [44, 49, 87]. Such work provides information on how one or a limited set of *a priori* regions are functionally related to other areas but does not reveal how all regions are interrelated, which was the goal of the present work. A few studies using optical imaging have gone beyond seed-based analysis; however, the number of identified networks in these studies was limited. For example, Vanni and colleagues [74] investigated cortical networks in GCaMP6 mice and reported three networks based on slow temporal frequencies (< 1 *Hz*) and two based on faster temporal frequencies (3 *Hz*) (see also [46]). Here, when we decomposed the cortex into three networks, we observed visual (OC-2) and somatomotor (OC-3) networks and a network that overlapped with territories possibly linked to a “default network” (OC-1) [13, 14, 18, 28] (Figure 2A). At this coarse scale, our results agreed with Vanni et al. [74] and other seed-based approaches [13, 20, 44, 49]. Importantly, we sought to determine functional organization at finer spatial levels too. With seven networks, we still observed visual (OC-2) and somatomotor (OC-3) networks, now together with a finer decomposition of other cortical systems (Figure 2C). Overall, our analyses reproduced previous observations at a coarse scale but characterized a more fine-grained decomposition of cortical organization.

The concordance between BOLD and Ca^2+^ networks was further evaluated quantitatively. When all regions were considered together, BOLD, 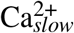, and 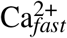 similarity was high with three networks (cosine similarity of ~ 0.85), and intermediate with either a seven-(0.7 – 0.8) or twenty-network (~ 0.65) decomposition. Similarity between 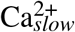 and 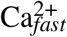 was high for all subdivisions (0.8 – 0.9) (Figure 2F). In the case of seven networks, we investigated pairwise network similarity (Figure 2D and E). Whereas some networks were similar across all comparisons (OC-1 to OC-4), others were not. OC-5 similarity between BOLD and 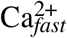 was very low (~ 0.6), while for OC-7, BOLD and either 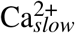 or 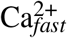 were “inversely correlated” (~ 0.26; i.e., largely spatially segregated).

We tested the hypothesis that BOLD networks are more similar to 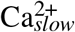 than 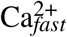 networks, which was motivated by the assertion that BOLD captures processes in the low-frequency regime [40, 45, 47, 49, 66–68]. In broad terms, considering all networks together, the hypothesis was confirmed for three, seven, and twenty networks, and when the number of ROIs and functional connectivity matrix thresholds were varied (Figure S2 and Figure S3). However, this conclusion should be qualified by the finding that 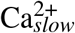 and 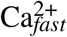 network results were in close agreement. In other words, for the temporal frequency ranges investigated here, Ca^2+^–based functional systems at low vs. high temporal frequencies were quite similar. Thus, linking the functional organization obtained with BOLD to 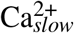 signals is not fully supported by our findings.

For the seven-network solution, OC-5 and OC-6 BOLD and 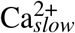 were similar (cosine similarity > 0.8) but deviated from the corresponding 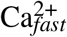 networks (~ 0.6). In line with this finding, 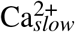 and 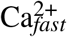 similarity was on the lower end (0.5 – 0.7). The anatomical areas that make up these networks may lend some insight into this finding. For example, OC-5 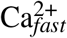 captured portions of the supplemental somatosensory area and barrel cortex, in addition to frontal areas including portions of the anterior lateral motor area. Neurons in the supplemental somatosensory area have long-range projections to the anterior lateral motor area [88], and their long-range coordination is thought to mediate tactile decision making [88, 89]. It is possible that higher temporal frequency Ca^2+^ signals are a signature of this circuit and that this activity is not well captured by the BOLD signal.

Overall, the proposal that different bands capture distinct neurophysiological properties [69] was not supported by our findings of the organization of large-scale systems. Instead of a “universal” relationship, we uncovered three distinct scenarios. (1) Low and high frequency Ca^2+^ signals manifested networks that were also recovered by BOLD (OC-1 to OC-4). (2) Networks that were somewhat separable by frequency band; specifically, low but not high frequency Ca^2+^ networks better matched their BOLD counterparts (OC-5 and OC-6). (3) Networks that were dissimilar across modalities regardless of Ca^2+^ temporal frequency (OC-7).

### Overlapping organization strongly represented in mouse BOLD and Ca^2+^ data suggests neural origins

Work in humans provides evidence for overlapping network organization based on BOLD [7, 58–60]. By some estimates, more than 40% of the human cortical surface participates in multiple networks, with some networks (e.g., default and dorsal attention) being comprised of nodes exhibiting > 90% overlap [59] (see also [7, 58]). However, as BOLD signals are indirect measures of neural activity, it is possible that such organization, at least in part, is driven by non-neuronal contributions, such as astrocytes [90] or vascular effects [31, 32, 35, 36].

To address this question in the mouse cortex, we quantified mixed membership in BOLD and wide-field Ca^2+^ imaging data. First, we investigated the distribution of membership values. A purely disjoint organization would be characterized by a skewed distribution where nearly all regions would display high membership values (indicating near-exclusive network participation). Simulations of synthetic disjoint graphs (i.e., zero overlap) showed that nodes have membership values > 0.8 (Figure S5). This was in stark contrast to our data, where even the networks with the largest numbers of “exclusive regions” (i.e., participating mostly in one network) exhibited considerable overlap. Specifically, across the three analysis conditions (BOLD, 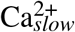, 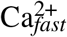), most networks had < 40% of regions in this high membership range; networks with the most disjoint organization were OC-7 (retrosplenial; 55% of regions were “exclusive”), OC-2 (visual; 60%), and OC-3 (somatomotor; 75%), which were observed with 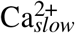 signal.

Further quantification of overlap was obtained by establishing the number of regions with membership values statistically greater than 1/7 (in the seven-network condition; Figure 4A). Using this definition, 55% of regions exhibited overlap for BOLD, 44% for 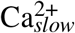, and 44% for 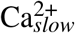 (Figure 4B and C). Thus, although BOLD networks exhibited the strongest overlap, both 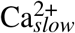 and 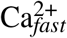 networks were also substantially overlapping. Further, several networks were highly overlapping across all three conditions (BOLD, 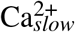, 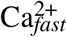); for example, OC-1 and OC-4 showed > 80% overlap (Figure 4D).

Together, these results show that network overlap is a prominent property of the functional organization of the mouse cortex for both sources of signal contrast. Critically, although the extent of network overlap was largest for BOLD, it was also pronounced in Ca^2+^ signals regardless of temporal frequency. These results lend support for the existence of the considerable overlapping organization in the human brain discovered with BOLD [7, 58–60].

### Network membership diversity is correlated across modalities but with notable differences

To characterize region-level properties of cortical organization, we quantified membership diversity (via entropy), which captures the extent to which regions affiliate with multiple networks. As expected, Ca^2+^ entropy maps showed lower diversity in sensorimotor regions than areas that have been previously implicated in multiple processes (Figure 5B). In particular, the posterior parietal cortex, which includes some higher visual areas, was among the regions with the highest diversity, consistent with the fact that it is widely anatomically connected [91] and involved in multiple functions (including navigation and decision making [92]).

Surprisingly, despite showing a positive correlation with the Ca^2+^ results (Figure 5C), the membership diversity observed with BOLD showed some particular results. For example, in contrast with Ca^2+^ data, the posterior parietal cortex did not exhibit high membership diversity with BOLD. In addition, somatosensory areas such as SSp-tr and SSp-ll (associated with body trunk and lower limbs) exhibited high diversity based on BOLD data, which was unexpected given that these regions are not known to be functionally diverse; and again, the Ca^2+^ data produced the anticipated low diversity outcome. Overall, these discrepancies do not disrupt a positive correlation between BOLD and Ca^2+^ entropies but raise questions about the extent to which the two imaging techniques are capturing the same phenomena.

### Character of network overlap is robustly different across imaging modalities

Brain regions with high network diversity participate in multiple networks. Bertolero et al. [65] referred to such regions as members of a “diverse club” and suggested that they play a more central role in the integration of information in comparison to “rich club” regions (i.e., those with high degree). Here, to probe this issue, we determined the relationship between entropy and degree across regions. For BOLD, membership diversity and degree were inversely correlated, a pattern indicative of what we called “sparse overlap” (regions at sectors of overlap are not highly functionally connected; Figure 7), which has also been observed in human BOLD [64, 65]. In contrast, Ca^2+^ data exhibited a positive correlation (that was more pronounced for 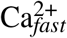), suggestive of more “dense overlap” (regions at sectors of overlap are highly functionally connected). To probe the robustness of this result, we computed node degree across functional connectivity thresholds, number of ROIs, as well as data preprocessing choices, and confirmed that the overall pattern persisted under all conditions (Figure S7). Together, these results revealed that BOLD and Ca^2+^ capture distinct forms of overlapping network organization. Following the proposal by Bertolero et al. [65] that the “diverse club” reveals regions that are more truly integrative in a network, we suggest that the Ca^2+^ results uncovered territories of network overlap corresponding to regions with stronger roles in signal integration and distribution. Future studies are needed to clarify why the organization observed with BOLD signals was different.

### Study considerations and future work

Due to the challenges presented by imaging mice while awake, including animal head motion and stress [93], we studied lightly anesthetized animals. Head motion systematically alters the correlation structure across brain regions [94, 95]. The effects of anesthesia on brain activity and neurovascular coupling are complex and may depend on brain region, anesthetic agent, and dose [87, 96–98]. However, the majority of human BOLD is collected from awake subjects. Thus, future studies will need to evaluate how functional networks, and their properties, differ between awake and anesthetized animals.

Processing and analyzing multimodal data entails making several parameter choices that could affect outcome measures. In particular, network overlap could be inflated by spatial misalignment. We took great care in co-registering animals by using the Allen Institute’s histological volume template and an optimized parameter set from the Advanced Normalization Tools package [99]. Further, issues of misalignment were considerably reduced by estimating network measures at the level of runs and combining values subsequently. Thus, modest misalignments after registration did not inflate the overall evaluation of overlap (see Methods). In addition, the quantification of membership strength was applied to values that were thresholded based on statistical significance. We also used relatively sparse graphs (15% density in the main text), such that only the strongest correlations were considered; further analyses that quantified the extent of overlap considered only membership values that statistically exceeded 1/7 (for the seven-network solution). We also probed the effects of parameter changes, including initial density of the correlation matrix, coarseness of cortical parcellation, and number of networks. Across all conditions, the results were qualitatively robust.

Although the findings reported in the main text were at the group level, our extensively sampled dataset allowed network organization to be determined at the level of the individual (Figure S4), lending considerable strength to our group-level findings, and underscoring the translational potential of our approach [100]. We plan to explore individualized network properties in subsequent work. Finally, because our multimodal data were collected simultaneously, the same ground truth brain activity underlies our results, such that moment-to-moment differences in spontaneous brain activity did not drive our findings.

### Conclusions

We employed novel simultaneous wide-field Ca^2+^ imaging and BOLD in a highly sampled group of mice expressing GCaMP6f in excitatory neurons to determine the relationship between large-scale networks discovered by the two techniques. Our findings demonstrated that (1) most BOLD networks were detected via Ca^2+^ signals. (2) Considerable overlapping network organization was recovered by both modalities. (3) The large-scale functional organization determined by Ca^2+^ signals at low temporal frequencies (0.01 – 0.5*Hz*), relative to high frequencies (0.5 – 5*Hz*), were more similar to those recovered with BOLD; yet, qualitative differences were also observed. (4) Key differences were uncovered between the two modalities in the spatial distribution of membership diversity and the relationship between region entropy (i.e., network affiliation diversity) and degree. These findings uncovered a distinct overlapping network phenotype across modalities. In sum, this work revealed that the mouse cortex is functionally organized in terms of overlapping large-scale networks that are observed with BOLD, lending fundamental support for the neural basis of such property, which is also observed in human subjects. The robust differences which were uncovered demonstrate that Ca^2+^ and BOLD also capture some complementary features of brain organization. Future work exploring these commonalities and differences, using simultaneous multimodal acquisition presented here, promises to help uncover how large-scale networks are supported by underlying brain signals in health and disease.

## Acknowledgements

We would like to thank all members of the Multiscale Imaging and Spontaneous Activity in Cortex (MISAC) collaboration at Yale University. We thank P. Brown for valuable input on the design and building of the telecentric lens holder. We thank J. Greenwood and the Neurotechnology Core at Yale University for modifying and building optics associated with the telecentric lens. We thank A. DeSimone, P. Brown, and the Yale School of Medicine electronics and machine shop. This work was supported by funding from NIH R01 MH111424 (R.T.C., M.C., and F.H.), U01 N2094358 (M.C. and R.T.C.), as well as 1RF1NS130069 and R21AG075778 (EMRL). H. Vafaii’s contribution to this research was supported in part by NSF award DGE-1632976.

## Materials and methods

### Experimental model and subject details

All procedures were performed in accordance with the Yale Institutional Animal Care and Use Committee and are in agreement with the National Institute of Health Guide for the Care and Use of Laboratory Animals. All surgeries were performed under anesthesia.

#### Animals

Mice were group housed on a 12-hour light/dark cycle. Food and water were available *ad libitum*. Cages were individually ventilated. Animals were 6-8 weeks old, 25-30g, at the time of the first imaging session. Animals (SLC, Slc17a7-cre/Camk2*α*-tTA/TITL-GCaMP6f also known as Slc17a7-cre/Camk2*α*-tTA/Ai93) were generated from parent 1 (Slc17a7-IRES2-Cre-D) and parent 2 (Ai93(TITL-GCaMP6f)-D;CaMK2a-tTA). Both were on a C57BL/6J background. To generate these animals, male CRE mice were selected from the offspring of parents with different genotypes, which is necessary to avoid leaking of CRE expression. Animals were originally purchased from Jackson labs.

#### Head-plate surgery

All mice underwent a minimally invasive surgical procedure enabling permanent optical access to the cortical surface (previously described here: [40]). Briefly, mice were anesthetized with 5% isoflurane (70/30 medical air/O_2_) and head-fixed in a stereotaxic frame (KOPF). After immobilization, isoflurane was reduced to 2%. Paralube was applied to the eyes to prevent dryness, meloxicam (2 mg/kg body weight) was administered subcutaneously, and bupivacaine (0.1%) was injected under the scalp (incision site). Hair was removed (NairTM) from the surgical site and the scalp was washed with betadine followed by ethanol 70% (three times). The scalp was removed along with the soft tissue overlying the skull and the upper portion of the neck muscle. Exposed skull tissue was cleaned and dried. Antibiotic powder (Neo-Predef) was applied to the incision site, and isoflurane was further reduced to 1.5%. Skull-thinning of the frontal and parietal skull plates was performed using a hand-held drill (FST, tip diameter: 1.4 and 0.7 mm). Superglue (Loctite) was applied to the exposed skull, followed by transparent dental cement (C&B Metabond^®^, Parkell) once the glue dried. A custom in-house-built head plate was affixed using dental cement. The head-plate was composed of a double-dovetail plastic frame (acrylonitrile butadiene styrene plastic, TAZ-5 printer, 0.35 *mm* nozzle, Lulzbot) and a hand-cut microscope slide designed to match the size and shape of the mouse skull. Mice were allotted at least 7 days to recover from head-plate implant surgery before undergoing the first of three multimodal imaging sessions.

### multimodal image acquisition

All mice, *N* = 10, underwent 3 multimodal imaging sessions with a minimum of 7 days between acquisitions. All animals underwent all imaging sessions. None were excluded prior to the study end-point. Data exclusion (based on motion etc.) is described below. During each acquisition, we simultaneously acquired fMRI-BOLD and wide-field Ca^2+^ imaging data using a custom apparatus and imaging protocol we have described previously [40]. Functional MRI data were acquired on an 11.7 *T* system (Bruker, Billerica, MA), using ParaVision version 6.0.1 software. During each imaging session, 4 functional resting-state runs (10 min each) were acquired. In addition, 3 runs (10 mins each) of unilateral light stimulation data were acquired. These data are not used in the present study. Structural MRI data were acquired to allow both multimodal registration and registration to a common space. Mice were scanned while lightly anesthetized (0.5 – 0.75% isoflurane in 30/70 O_2_/medical air) and freely breathing. Body temperature was monitored (Neoptix fiber) and maintained with a circulating water bath.

#### Functional MRI

We employed a gradient-echo, echo-planar-imaging sequence with a 1.0 sec repetition time (TR) and 9.1 ms echo time (TE). Isotropic data (0.4 *mm* × 0.4 *mm* × 0.4 *mm*) were acquired along 28 slices providing near whole-brain coverage.

#### Structural MRI

We acquired 4 structural images for multimodal data registration and registration to a common space. (1) A multi-spin-multi-echo (MSME) image sharing the same FOV as the fMRI data, with a TR/TE of 2500/20 *ms*, 28 slices, two averages, and a resolution of 0.1 *mm* × 0.1 *mm* × 0.4 *mm*. (2) A whole-brain isotropic (0.2 *mm* × 0.2 *mm* × 0.2 *mm*) 3D MSME image with a TR/TE of 5500/20 *ms*, 78 slices, and two averages. (3) A fast-low-angle-shot (FLASH) time-of-flight (TOF) angiogram with a TR/TE of 130/4 *ms*, resolution of 0.05 *mm* × 0.05 *mm* × 0.05 *mm* and FOV of 2.0 *cm* × 1.0 *cm* × 2.5 *cm* (positioned to capture the cortical surface). (4) A FLASH image of the angiogram FOV, including four averages, with a TR/TE of 61/7 *ms*, and resolution of 0.1 *mm* × 0.1 *mm* × 0.1 *mm*.

#### Wide-field fluorescence Ca^2+^ imaging

Data were recorded using CamWare version 3.17 at an effective rate of 10 *Hz*. To enable frame-by-frame background correction, cyan (470/24, Ca^2+^-sensitive) and violet (395/25, Ca^2+^-insensitive) illumination (controlled by an LLE 7Ch Controller from Lumencor) were interleaved at a rate of 20 *Hz*. The exposure time for each channel (violet and cyan) was 40 *ms* to avoid artifacts caused by the rolling shutter refreshing. Thus, the sequence was: 10 *ms* blank, 40 *ms* violet, 10 *ms* blank, 40 *ms* cyan, and so on. The custom-built optical components used for in-scanner wide-field Ca^2+^ imaging have been described previously [40].

### Image preprocessing

#### multimodal data registration

All steps were executed using tools in BioImage Suite (BIS) specifically designed for this purpose [40]. For each animal, and each imaging session, the MR angiogram was masked and used to generate a view that recapitulates what the cortical surface would look like in 2D from above. This treatment of the data highlights the vascular architecture on the surface of the brain (notably the projections of the middle cerebral arteries, MCA) which are also visible in the static wide-field Ca^2+^ imaging data. Using these and other anatomical landmarks, we generated a linear transform that aligns the MR and wide-field Ca^2+^ imaging data. The same static wide-field image was used as a reference for correcting motion in the time series. To register the anatomical and functional MRI data, linear transforms were generated and then concatenated before being applied.

Data were registered to a reference space (CCFv3, [70]) using isotropic whole-brain MSME images via affine followed by non-linear registration (ANTS, Advanced normalization tools; [99]). The histological volume in CCFv3 was used because of a better contrast match with MRI images. The goodness of fit was quantified using mutual information and a hemispheric symmetry score that captured the bilateral symmetry of major brain structures. A large combination of registration hyperparameters was explored, and the top 10 fits per animal were selected. The best transformation out of this pool was selected for each animal by visual inspection.

#### Fluorescence Ca^2+^ imaging data preprocessing

We have previously described all steps in this pipeline [72]. Briefly, the raw signal was split between GCaMP-sensitive and GCaMP-insensitive imaging frames. Spatial smoothing with a large kernel (16-pixel kernel, median filter) was applied to reduce and/or remove focal artifacts (e.g., dust or dead pixels from broken fibers). Focal artifacts do not move with the subject and can bias motion correction. Motion correction parameters were estimated on these data using normalized mutual information. Rigid image registration was performed between each imaging frame in the time series and the reference frame. Registration parameters were saved, and the large kernel-smoothed images were discarded. Modest spatial smoothing (4-pixel kernel, median filter) was applied to the raw data, and these data were motion corrected by applying the parameters estimated in the previous step. Data were down-sampled by a factor of two in both spatial dimensions, which yielded a per pixel spatial resolution of 50 × 50 *μm*^2^ (original was 25 × 25 *μm*^2^). Photo bleach correction was applied to reduce the exponential decay in fluorescence at the onset of imaging [101]. The fluorophore-insensitive time series were regressed from the fluorophore-sensitive time series. The first 50 seconds of data were discarded due to the persistent effects of photobleaching in the Ca^2+^ data. Data were band-pass filtered (3rd order Butterworth) between [0.01 – 0.5] *Hz* 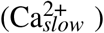, and high-pass filtered between [0.5 – 5.0] *Hz* 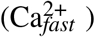, and 15 time points (1.5s of data) were discarded from both beginning and the end of the time series to avoid filtering-related edge artifacts [95].

#### RABIES fMRI data preprocessing

For fMRI preprocessing, we used RABIES (Rodent automated BOLD improvement of EPI sequences) v0.4.2 [71]. We applied functional inhomogeneity correction N3 (nonparametric nonuniform intensity normalization; [102, 103]), motion correction (ANTS, Advanced normalization tools; [99]), and slice time correction, all in native space. A within-dataset common space was created by nonlinearly registering and averaging the isotropic MSME anatomical images (one from each mouse at each session), which was registered to the Allen CCFv3 reference space using a nonlinear transformation (see above).

For each run, fMRI data were motion corrected and averaged to create a representative mean image. Each frame in the time series was registered to this reference. To move the fMRI data to the common space, the representative mean image was registered to the isotropic structural MSME image acquired during the same imaging session. This procedure minimizes the effects of distortions caused by susceptibility artifacts [104]. Then, the three transforms — (1) representative mean to individual mouse/session isotropic MSME image, (2) individual mouse/session isotropic MSME image to within-dataset common space, and (3) within-dataset common space to out-of-sample common space — were concatenated and applied to the fMRI data. Functional data (0.4mm isotropic) were upsampled to match anatomical MR image resolution (0.2mm isotropic). Registration performance was visually inspected and verified for all sessions. Motion was regressed (6 parameters). Data were high-pass filtered (3rd order Butterworth) between [0.01 – 0.5] *Hz*, and 15 time points (15s of data) were discarded from both the beginning and the end of the time series to avoid filtering-related edge artifacts. Average white matter and ventricle time courses were regressed.

#### Frame censoring

Data were scrubbed for motion using a conservative 0.1 *mm* threshold. High-motion frames were selected based on estimates from the fMRI time series and applied to both fMRI and Ca^2+^ data. Runs were removed from the data pool if half of the imaging frames exceed this threshold for a given run. In this dataset, 2 runs (or ~ 1.7% of all runs) were removed for this reason. Additionally, 2 more runs were removed because they did not pass our quality control criteria.

### Parcellating the cortex into columnar regions of interest (ROI)

To create regions of interest (ROIs), we employed the Allen CCFv3 (2017) reference space [70] and used their anatomical delineations as our initial choice of ROIs. However, this led to poor performance (see supplemental discussion). Here, we introduce a novel spatially homogeneous parcellation of the mouse cortex that can be adopted for both 3D fMRI and 2D wide-field Ca^2+^ imaging data.

The procedure worked as follows. (1) We generated a cortical flatmap within the CCFv3 space using code published in ref. [105](https://github.com/AllenInstitute/mouse_connectivity_models). (2) We subdivided the left hemisphere into *N* regions via *k*-means clustering applied to pixel coordinates (for most analyses reported, *N* = 512). The right hemisphere was obtained by simple mirror-reversal to obtain a total of 2*N* regions. (3) Depth was added to the ROIs to obtain column-shaped regions. To do so, a path was generated by following streamlines normal to the surface descending in the direction of white matter (streamline paths were available at 10 *μm* resolution in CCFv3; see Figure 3F in [70]). Here, we chose ROI depths so that we included potential signals from approximate layers 1 to 4 (layer masks were obtained from CCFv3). Evidence from wide-field Ca^2+^ imaging suggests that signals originate from superficial layers but can extend into the cortex to some extent [51, 81, 106–108]. (4) Finally, ROIs were downsampled from 10 *μm* to 100 *μm* resolution. See Figure 1C.

After co-registration, ROIs were transformed from the CCFv3 space into each individual’s 3D and 2D anatomical spaces (see above). On average, ROIs had a size of 8 ± 3 voxels (3D, fMRI) and 48 ± 20 pixels (2D, Ca^2+^) in individual spaces (mean ± standard deviation).

### Functional network construction

Time series data were extracted and averaged from all voxels/pixels within an ROI in native space to generate a representative time series per ROI. For each modality, for each run, an adjacency matrix was calculated by applying Pearson correlation to time series data to each ROI pair. Next, we binarized the adjacency matrices by rank ordering the connection weights and maintaining the top 15%; thus, after binarization, the resulting graphs had a fixed density of *d* = 15% across runs and modalities. This approach aims to keep the density of links fixed across individuals and runs and better preserves network properties compared to absolute thresholding [109]. To establish the robustness of our results to threshold values, we also tested values of 10% to 25% in 5% increments.

### Finding overlapping communities

Overlapping network analysis was applied by using SVINET, a mixed-membership stochastic blockmodel algorithm [61, 110], which has been previously applied to human fMRI data by us [58] and other groups [60]. SVINET models the observed graph within a latent variable framework by assuming that the existence (or non-existence) of links between pairs of nodes can be explained by their latent community memberships. For binary adjacency matrix *A* and membership matrix *π*, the model assumes the conditional probability of a connection as follows

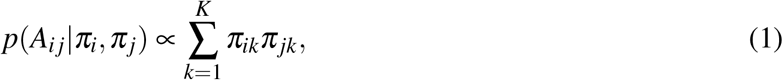

where *K* is the number of communities, and *A_ij_* = 1 if nodes *i* and *j* are connected and 0 otherwise. Intuitively, pairs of nodes are more likely to be connected if they belong to the same network or to (possibly several) overlapping networks. More formally, SVINET assumes the following generative process

1. For each node, draw community memberships *π_i_* ~ Dirichlet(*α*)
2. For each pair of nodes *i* and *j*:

a. Draw community indicator *z*_*i*→*j*_ ~ *π_i_*
b. Draw community indicator *z*_*i*←*j*_ ~ *π_j_*
c. Assign link between *i* and *j* if *z*_*i*→*j*_ = *z*_*i*←*j*_.

Model parameters *α* are fit using stochastic gradient ascent [111, 112]. The algorithm was applied to data from each run using 500 different random seeds. Results across seeds were combined to yield a final consensus for a run.

#### Aligning community results

Networks were identified in random order due to the stochastic nature of our algorithm. Maximum cosine similarity of the cluster centroids was used to match networks across calculations (runs or random seeds). For each run and random seed, membership vectors for all random seeds were submitted to *k*-means clustering (sklearn.cluster.KMeans) to determine *K* clusters (e.g., *K* = 7 for analyses with 7 networks). The similarity between the resulting cluster centroids was then established via cosine similarity, and the networks matched based on maximum similarity. Formally, the outcome was membership matrix *π* (Figure 1E).

### Group results

Crucially, all measures were computed at the run level first before combining at the group level.

#### Membership matrices

This is what’s visualized in Figure 2A and C, Figure S1 and Figure S2A and C.

#### Thresholding membership values

To enhance the robustness of our estimates of network overlap, membership values were thresholded to zero if they did not pass a test rejecting the null hypothesis that the value was zero. After thresholding, the surviving membership values were rescaled to sum to 1. Thresholding was performed for each animal separately by performing a one-sample *t*-test and employing a false discovery rate of 5%. All results shown utilized this step, with the exception of figures which illustrate the spatial patterns of membership values (and do not estimate network overlap). Note that almost all (~ 99%) memberships that did not reach significance had values in the range [0,0.2].

#### Region functional diversity

Shannon entropy was applied to membership matrices for each run separately before averaging. That is, given a membership matrix *π* from a run, node entropies were computed to get an entropy estimate per node at the run level

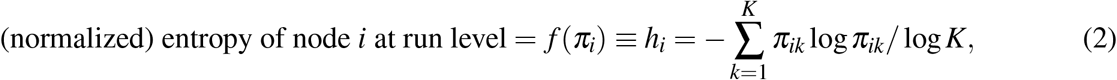

where *K* = 7 is the number of communities. Entropy values were combined by averaging over runs to get the group-level estimates. This is what’s visualized in Figure 5B. Similarly, group averages were used to calculate the correlations between modalities in Figure 5C.

#### Computing distributions

Similar to above, distributions were computed for each run separately before combining at the group level. For example, consider *h_i_* (entropy of node *i*) from a run. We computed percentage values using 20 bins of width 0.05 that covered the entire range of normalized entropy values [0,1]. We then averaged over run-level histogram values to get group-level estimates shown in Figure 5A. Other distributions were computed in an identical way. Specifically, 57 bins of size 5 were used for Figure 6A, and 4 bins of size 0.2 were used for Figure 3B.

### Statistical analysis

#### Hierarchical bootstrapping

Statistical results were performed at the group level by taking into consideration the hierarchical structure of the data (for each animal, runs within sessions), which can be naturally incorporated into computational bootstrapping to estimate variability respecting the organization of the data [113]. For each iteration (total of 1,000, 000), we sampled (with replacement) *N* = 10 animals, *N* = 3 sessions, and *N* = 4 runs, while guaranteeing sessions were yoked to the animal selected and runs were yoked to the session selected (Figure 1B). In this manner, the multiple runs were always from the same session, which originated from a specific animal. Overall, the procedure allowed us to estimate population-level variability based on the particular sample studied here. To estimate 95% confidence intervals, we used the bias-corrected and accelerated (BCa) method [114], which is particularly effective when relatively small sample sizes are considered (SciPy’s scipy.stats.bootstrap). See Figure 2E and F, Figure 3B, Figure 5C, Figure 6C, Figure 7B, Figure S2E and F, Figure S3, and Figure S6.

#### One-sample t-test

We used a one-sample *t*-test to define *belonging* in communities as a function of a threshold *μ*. The *t*-statistic for membership of node *i* in community *k* is given as

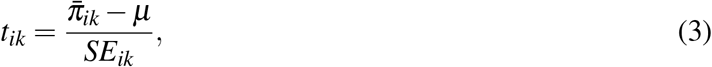

where 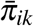 is the group averaged membership of node *i* in community *k*, and *SE_ik_* is the standard error estimated using hierarchical bootstrapping (see above). We calculated p values using *t*-statistics for all nodes and communities and declared a node a member of a community if its p-value reached significance *p* = 0.05. The results as a function of various *μ* are shown in Figure 4A. We applied Benjamini-Hochberg correction [115] using Python statsmodels’ implementation (statsmodels.stats.multitest.multipletests) to correct for multiple comparison.

#### LFR analysis

The following parameters need to be specified to generate a binary and overlapping LFR graph [84]. *N*, number of nodes; *k*, average degree; *μ*, topological mixing parameter; *t*_1_, minus exponent for the degree sequence; *t*_2_, minus exponent for the community size distribution; *C_min_*, minimum for the community sizes; *C_max_*, maximum for the community sizes; *ON*, number of overlapping nodes; *OM*, number of memberships of the overlapping nodes.

To match basic statistics of the real data with LFR graphs we set *N* = 542 (Figure 1D); and, for every run from each data modality, we calculated the average degree *k* and estimated *t*_1_ via an exponential fit to the degree distributions (scipy.stats). We set *t*_2_ = 0.1, *C_min_* = 0.05 × *N* ≈ 27, *C_max_* = 0.35 × *N* ≈ 190. For the fraction of overlapping nodes *ON*, we explored a wide range between 0 (disjoint) and 0.9 in incremental steps of 0.1. This yielded *ON* = 0 up to *ON* = 0.9 × *N* = 488. Finally, we used *OM* = 2 and 3. This results in a total of 20 LFR graphs per run, per data modality. We applied the community detection algorithm to LFR graphs in an identical way to the real data, but with fewer seeds (*N* = 10 compared to *N* = 500). The alignment procedure was performed in an identical way as described above. See Figure S5.

#### Permutation test

A paired permutation test was used to compare conditions in Figure 2E and F, Figure S2E and F, and Figure S3; and to perform a node-wise comparison across modalities in Figure 5D. We used SciPy’s implementation (scipy.stats.permutation_test) with *N* = 1,000,000 resamples. Holm–Bonferroni correction was applied to correct for multiple comparison [116].

## Supplemental information

### Supplemental discussion

#### Choosing number of communities

Clustering is in the eyes of the beholder (or the algorithm) [117]. Community detection is inherently ill-defined: algorithms do not find communities, what they do is “*define*” communities and later find them according to their definition. Here, we employed a mixed-membership stochastic blockmodel [73], where each community corresponds to a *latent functional role* (see [118] for an extensive review). Choosing a specific number of communities is thus equivalent to deciding how many functional roles (or clusters) best describe the observed graphs. In this sense, there is no “*true*” number of communities. Therefore, instead of focusing on a single result as a true decomposition, we explored network organization at different levels of granularity. We decided the number of communities empirically, using criteria such as bilateral symmetry. At the most coarse level, our *K* = 3 communities recapitulated previous seed-based reports in Ca^2+^ data and coarse fMRI-ICA findings. Our *K* = 7 decomposition was similar to previous fMRI-ICA reports [15, 20]. Overall, the highly bilateral nature of the resulting community structure (observed for up to *K* = 20 for Ca^2+^) and their similarity to previous reports increased our confidence about our results.

#### Thresholding the graphs

The algorithm used in this study requires binary graphs as its input [61], which necessitates choosing an edge-filtering threshold. To mitigate this, we employed an approach known as proportional thresholding which is known to perform better than alternatives such as absolute thresholding [109]. In addition, theoretical work has shown that network topology is highly robust against different thresholds [119]. Here we reported results at a graph density of *d* = 15%. To ensure the robustness of our findings to this arbitrary choice, we also explored densities going from *d* = 10% all the way up to *d* = 25% with incremental steps of 5%. Empirically, we found that our results were robust to the choice of edge density, as well as other hyperparameters. Overall, our data partially confirmed previous theoretical and simulation work that network topology is robust across a wide range of sparsity levels.

#### Region of interest (ROI) definition scheme

We started by using brain region masks from Allen Reference Atlas (ARA) as our initial choice of ROIs [70]. However, we observed some mismatch between functional parcellations and ARA parcellation. This is probably because ARA regions were delineated using various anatomical and structural criteria; but crucially, function was not one of them. In the present work, we introduced a new parcellation scheme illustrated in Figure 1C and D, which increased the robustness of our results. In conclusion, we found that spatially homogeneous ROIs worked well for the purpose of functional network construction, consistent with previous reports in humans [120].

Defining appropriate ROIs for our multimodal dataset was challenging because different modalities occupy spaces with different geometries. Namely, fMRI data is defined within a 3D volumetric space, while mesoscopic Ca^2+^ imaging data exists only on the 2D cortical surface. To address this, we started from the 2D space of cortical flatmap (Figure 1C; step I), which fits Ca^2+^ data well. We then added depth to obtain 3D volumetric ROIs (Figure 1C; step II), suitable for fMRI data. In the depth dimension, we included layers 1 to 4. Crucially, our goal was not to obtain layer-specific results (BOLD resolution was 0.4 *mm* isotropic). Instead, we wanted to consider depths of the cortex that most likely contribute to Ca^2+^ signal [51, 81, 106–108], which would render our network-level cross-modality comparisons more meaningful.

#### Comparing entropy to other measures of node diversity

Brain regions of high functional diversity are more likely to participate in multiple networks. This region-level property can be characterized using appropriate node centrality measures. Here, we took advantage of having access to continuous membership values and defined node entropy centrality (Figure 5). Another measure, *“participation coefficient”* [17, 64, 65, 85, 86], has also been commonly used to quantify a similar concept: a node’s participation coefficient measures how well-distributed its links are among different communities. Large participation coefficient indicates higher amount of link diversity, which could be potentially related to high membership diversity. To understand the relationship between the two measures, we visualized their spatial patterns and found that entropy and participation coefficient maps were largely in agreement (Figure S6). The node-wise correlation between entropy and participation coefficient was *r* = 0.70 ± 0.09, BOLD; *r* = 0.77 ± 0.12, 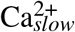; *r* = 0.47 ± 0.28, 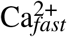.

It is worth noting that node entropy and participation coefficient are defined in very different ways. Entropy is computed from membership probability vectors within our overlapping framework; whereas, participation coefficient depends on how links are distributed across communities within a disjoint framework. Despite this, the two measures were highly correlated indicating that they probably capture similar underlying phenomena.

**Figure S1:**
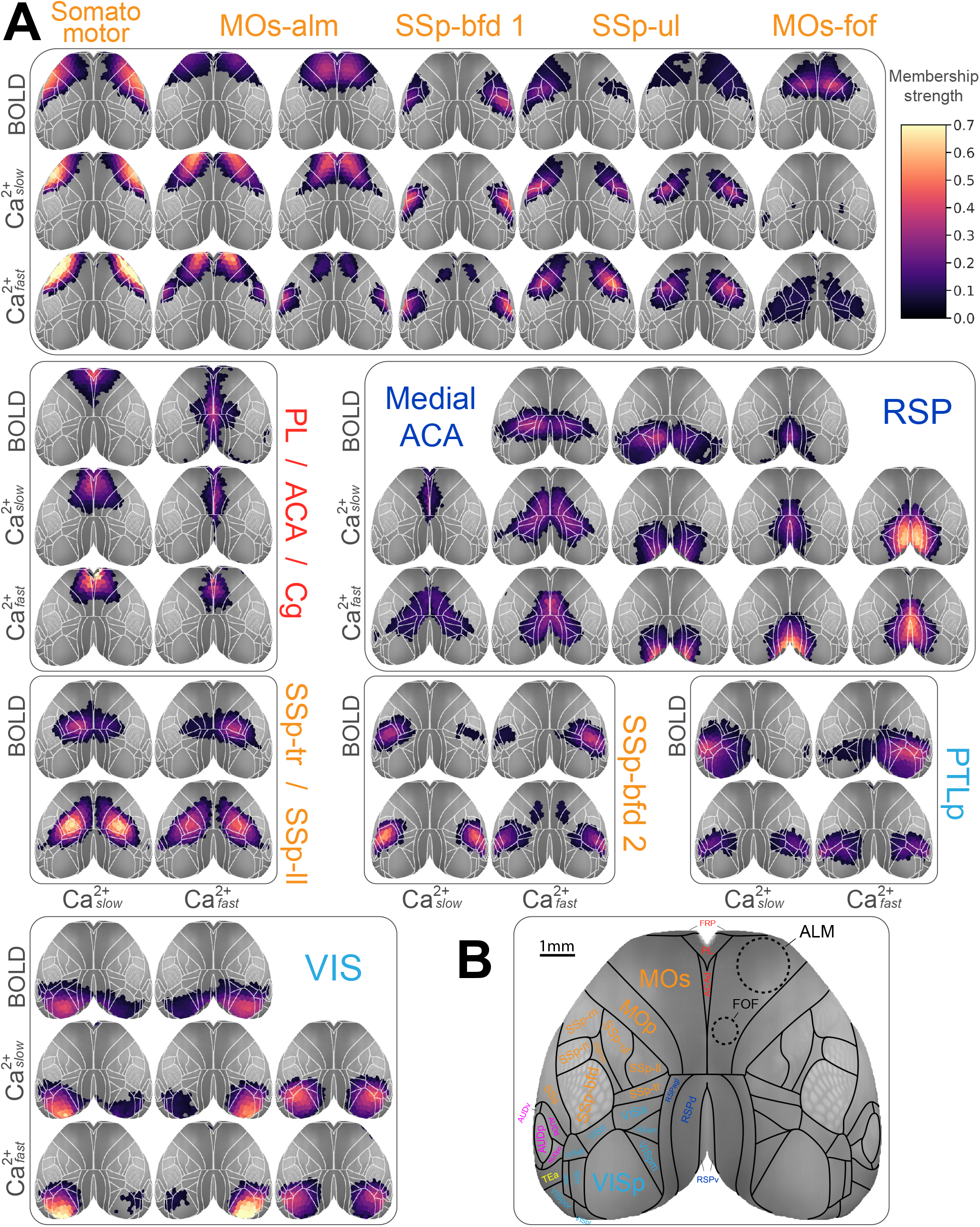
*K* = 20 decomposition. **(A)** Even at *K* = 20, most networks maintain their bilateral symmetry, especially for Ca^2+^. A network centered around FOF appears as its own separate network for BOLD, similar to the *K* = 7 solution (top-right). In contrast, this network did not appear separately for Ca^2+^, even at *K* = 20. Instead, FOF partially overlaps with a large medial Ca^2+^ network that spans parts of SSp-tr/ll. **(B)** Fine divisions of the cortical regions in Allen reference atlas, along with ALM and FOF (dashed lines). Label colors inspired from Figure 1F in ref. [121]. A complete list of abbreviations can be found in the original CCFv3 publication [70]. Compare with Figure 2 for *K* = 3 and *K* = 7 solutions.

**Figure S2:**
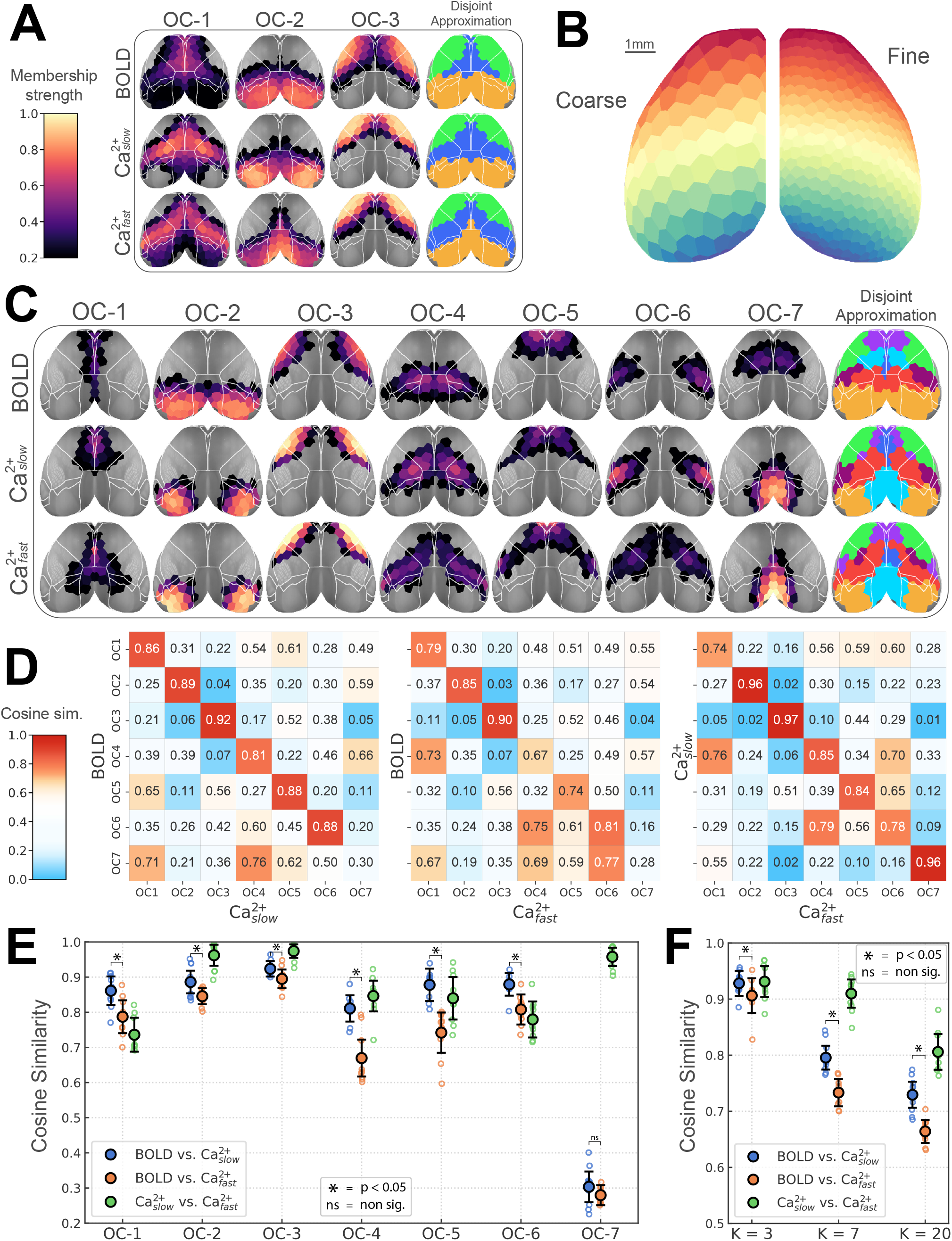
Network structure is robust to the choice of ROI granularity. **(A, C-F)** Similar to Figure 2. **(B)** Left, results with coarse ROIs are presented here; Right, fine ROIs were used for the main results. See also Figure S3.

**Figure S3:**
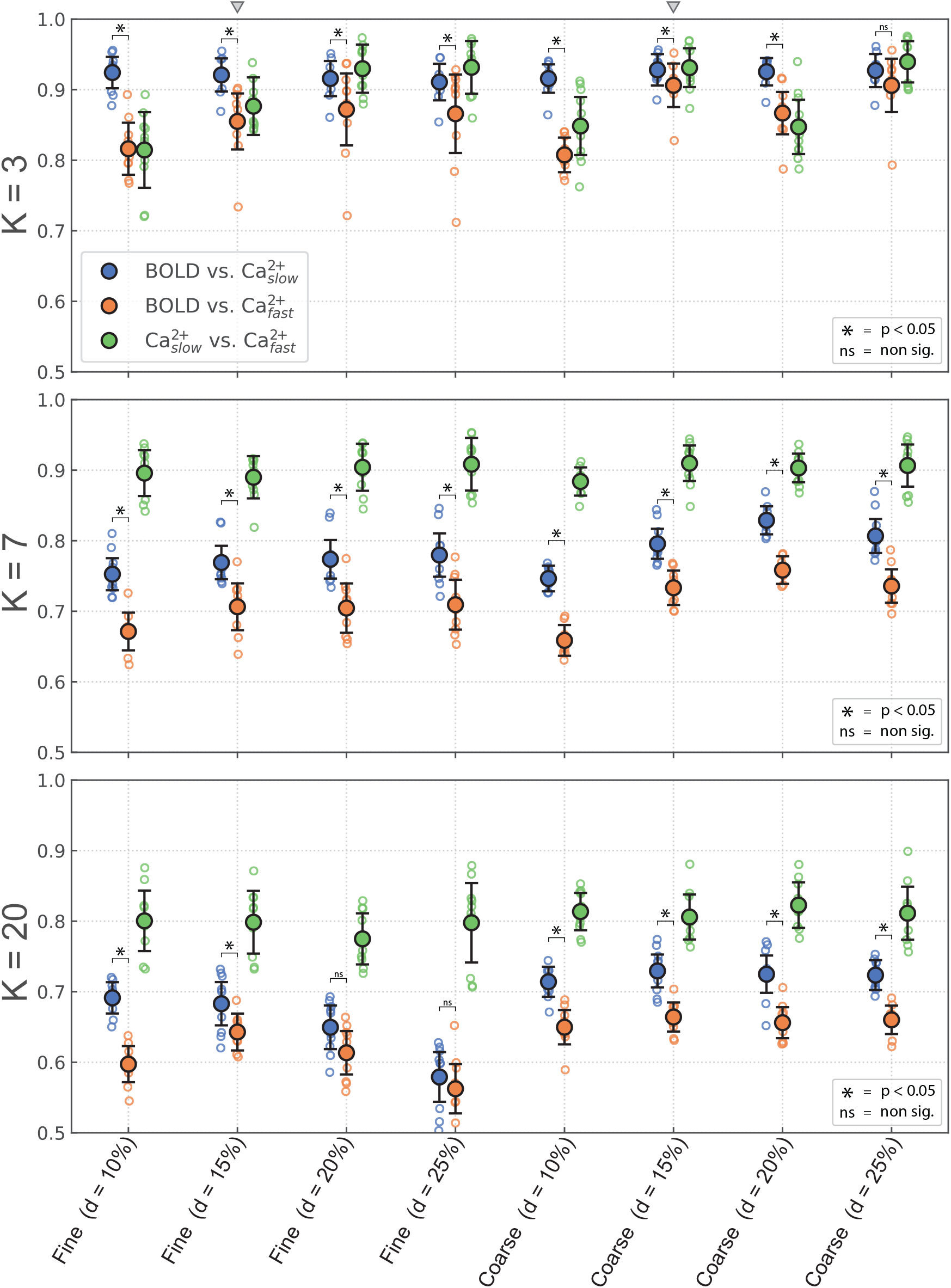
BOLD network organization is more similar to 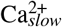 than it is to 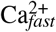. This is a robust finding in the present study, as it is reproduced over a large combinatorial space of analysis conditions. The y-axis is cosine similarity and the x-axis corresponds to different conditions. See Figure S2B for a visual comparison of coarse versus fine ROIs. *d* is graph density after edge-filtering is applied. Small triangles indicate our choices for the main results: *d* = 15%, fine ROIs. Permutation test, *p* < 0.05, Holm-Bonferroni corrected. Error bars show 95% confidence intervals, hierarchical bootstrap. See also Figure 2 and Figure S2.

**Figure S4:**
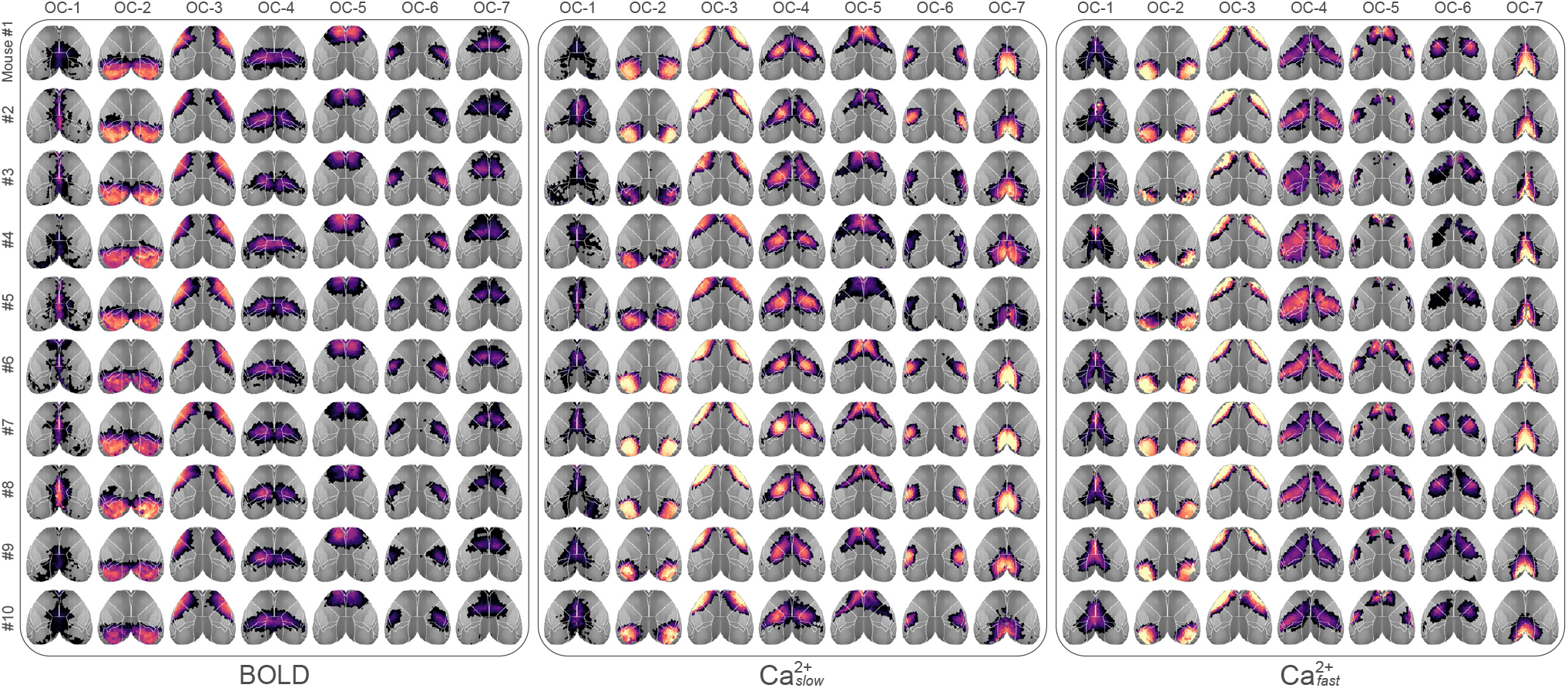
Overlapping networks at the level of individuals. Each mouse was highly sampled (Figure 1B), which allowed robust estimation of individualized networks. The high similarity of network structures across individuals is anticipated; however, there are still visible differences. Overall, the reproducibility of results at the individual level is notable and lends support to our group results. Compare with Figure 2C for group results.

**Figure S5:**
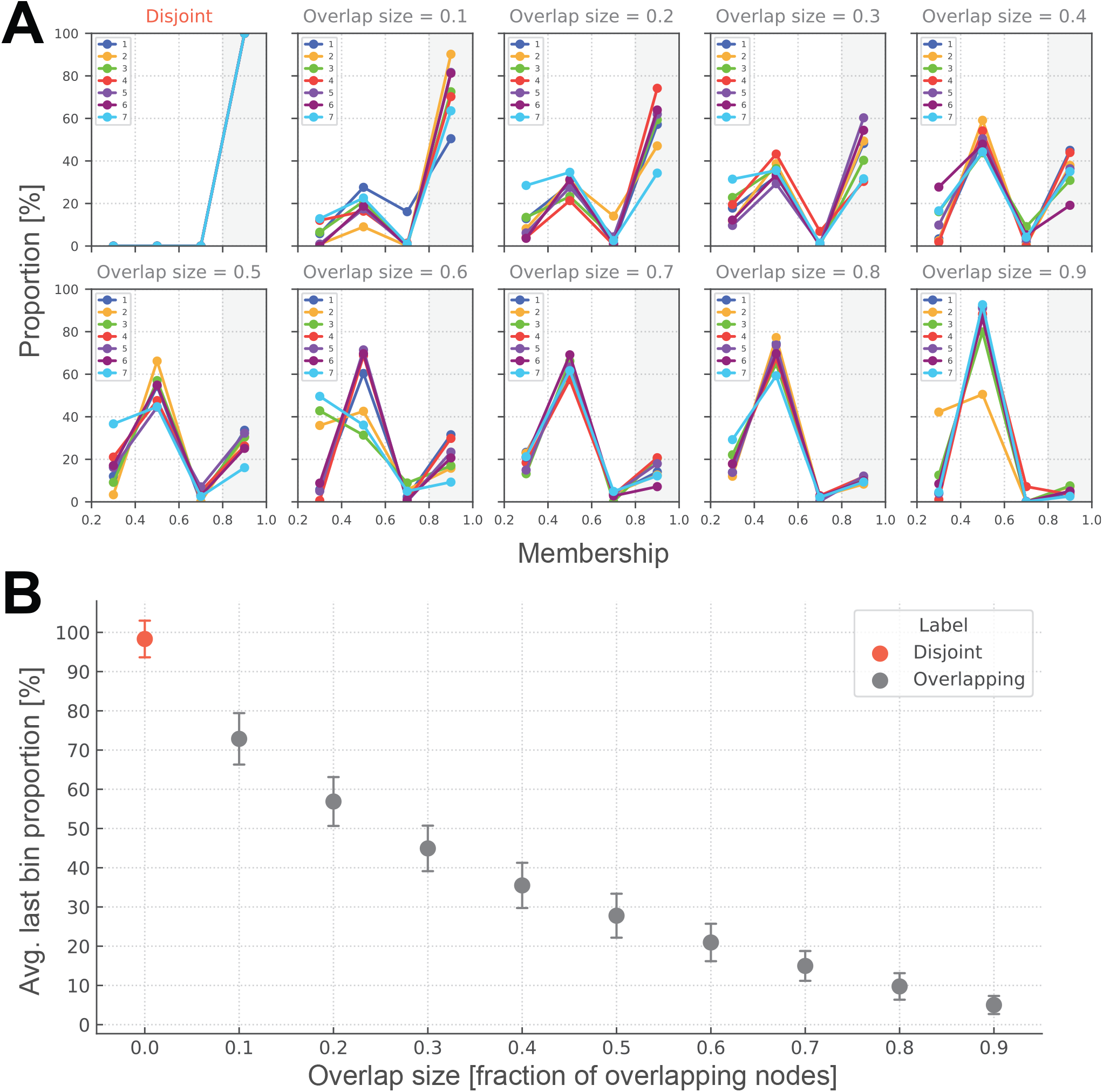
Verifying our analysis procedure using synthetic LFR graphs [84]. **(A)** To compute the distribution of node memberships, we divided the interval (0.2,1.0] into four bins of equal width. We chose this binning scheme because it allowed us to distinguish between disjoint and overlapping graphs: for disjoint graphs, the bin corresponding to strong membership values (i.e., last bin) had a proportion of 100%, and the proportions were 0% elsewhere. Overlap size, or fraction of overlapping nodes, is a tunable parameter in LFR graphs. Here we show how membership distributions change as a function of overlap size. **(B)** We averaged the last bin proportions across all communities to get a single statistic per graph. Its mean over thousands of simulated graphs is shown here. The average last bin proportion drops as overlap size increases, thus making it a reliable proxy for network overlap size. Error bars indicate standard deviation. Compare with Figure 3.

**Figure S6:**
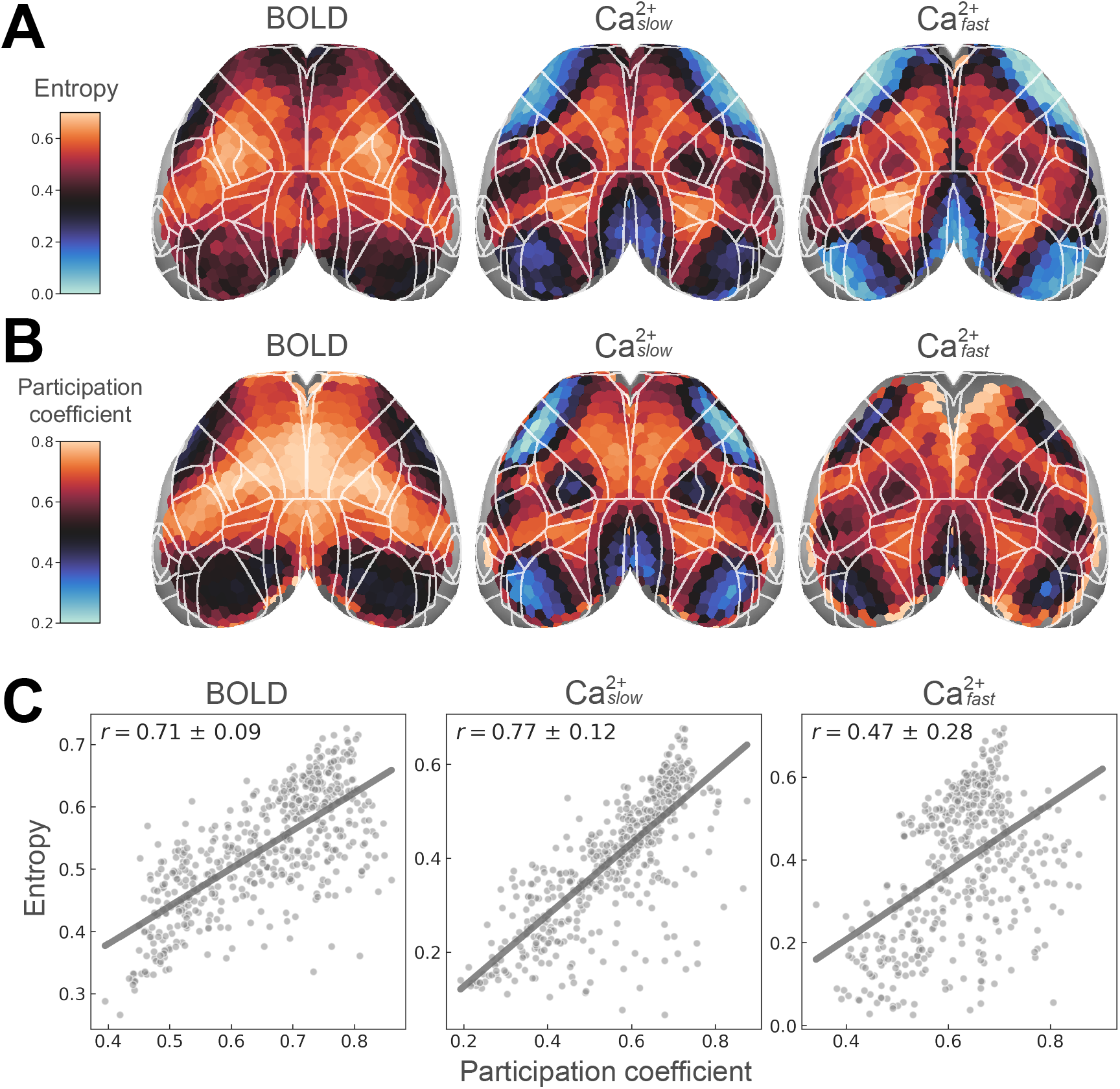
Entropy and participation coefficient uncover similar spatial patterns. **(A)** Here, we show entropy maps with actual values (compare with Figure 5B for the rank-ordered version). Note the positive shift in BOLD values. **(B)** Participation coefficient is commonly used to quantify how a node’s links are distributed across (disjoint) communities [17, 64, 65, 85, 86]. To compute participation coefficients, we used the disjoint approximation obtained from taking the maximum membership of a given node (see the last column in Figure 2C). Note that for low-degree nodes with degree < 7, participation coefficient estimates become unreliable. For example, see frontal regions in 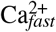. **(C)** The two centrality measures are correlated, suggesting that they capture similar underlying phenomena. The concordance between these measures is most clearly visible for 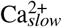, but also for BOLD to some extent.

**Figure S7:**
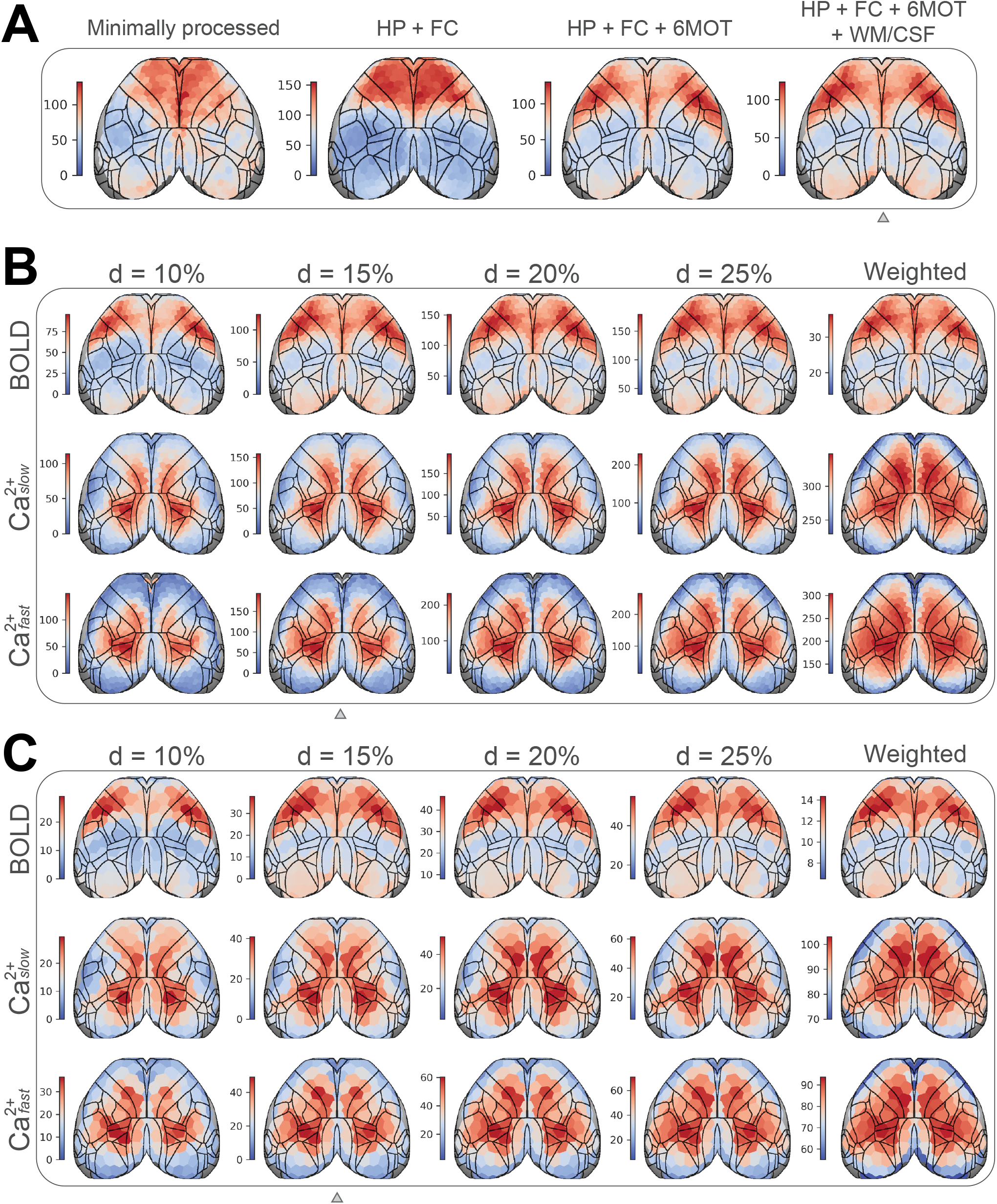
Dependence of degree centrality to preprocessing and analysis choices. **(A)** Spatial pattern of average node degree is somewhat altered depending on which BOLD preprocessing steps are used. Minimally processed, motion correction (rigid transformations) and detrending; HP, high-pass filtering (0.01 *Hz*); FC, frame censoring; 6MOT, motion regression (6 parameters); WM/CSF, average signal from white matter and ventricles regressed out. **(B)** Degree patterns are robust to the choice of edge filtering threshold. Difference thresholds result in different scales, but the spatial patterns remain relatively similar. **(C)** Similar to B but for a coarse parcellation (see Figure S2B). Small triangles indicate the pipeline and parameters used for the main results. Related to Figure 6.

